# The Kinesin-14 Tail: Dual microtubule binding domains drive spindle morphogenesis through tight microtubule cross-linking and robust sliding

**DOI:** 10.1101/2025.02.25.640188

**Authors:** Stephanie C. Ems-McClung, MacKenzie Cassity, Anjaly Prasannajith, Claire E. Walczak

**Author notes:** Department of Molecular Biology, Princeton University, Princeton, NJ 08540.

## Abstract

Proper spindle assembly requires the Kinesin-14 family of motors to organize microtubules (MTs) into the bipolar spindle by cross-linking and sliding anti-parallel and parallel MTs through their motor and tail domains. How they mediate these different activities is unclear. We identified two MT binding domains (MBD1 and MBD2) within the *Xenopus* Kinesin-14 XCTK2 tail and found that MBD1 MT affinity was weaker than MBD2. Comparable to full-length GFP-XCTK2 wild-type protein (GX-WT), GFP-XCTK2 containing the MBD1 mutations (GX-MBD1^mut^) stimulated spindle assembly, localized moderately on the spindle, and formed narrow spindles. In contrast, GX-MBD2^mut^ only partially stimulated spindle assembly, localized weakly on the spindle, and formed shorter spindles. Biochemical reconstitution of MT cross-linking and sliding demonstrated that GX-MBD2^mut^ slid anti-parallel MTs faster than GX-WT and GX-MBD1^mut^. However, GX-WT and GX-MBD1^mut^ statically cross-linked the majority of parallel MTs, whereas GX-MBD2^mut^ equally slid and statically cross-linked parallel MTs without affecting their sliding velocity. These results provide a mechanism by which the two different MT binding domains in the Kinesin-14 tail balance anti-parallel MT sliding velocity (MBD1) and tight parallel MT cross-linking (MBD2), which are important for spindle assembly and localization, and provide a basis for characterizing how molecular motors organize MTs within the spindle.

**Significance Statement:** Spindle assembly and organization utilize molecular motors that cross-link and slide anti-parallel and parallel microtubules. How individual motors moderate both active sliding and static cross-linking is not understood. Using biochemical reconstitution, the authors determined that the Kinesin-14 tail contains two independent microtubule binding domains. MBD1 with weaker microtubule binding facilitates faster anti-parallel microtubule sliding, whereas the stronger MBD2 mediates tight parallel microtubule cross-linking, which was important for spindle assembly. These findings provide a mechanism for how Kinesin-14s differentially control microtubule sliding and cross-linking and provide insight into how molecular motors can mediate the dynamic organization of microtubules in the spindle.

## Introduction

Spindles are composed of microtubules (MTs) that are organized into a bipolar array by the action of MT-associated proteins and molecular motors, including the kinesin superfamily (Valdez *et al*., 2023). Most MT minus ends are focused at the spindle poles, resulting in a parallel arrangement in each half spindle. In the middle of the spindle, the plus ends of MTs extending from each of the poles overlap at the equator creating an anti-parallel arrangement. Molecular motors that set up this array convert the chemical energy of ATP into force production for plus- or minus-end directed motility and/or for the regulation of MT dynamics (Dujardin and Vallee, 2002; Canty *et al*., 2021; Gassmann, 2023; Yildiz, 2024). Motile kinesins can be either plus-end directed or minus-end directed motors, and during spindle assembly move chromosomes, transport MTs, and/or regulate MT dynamics (Yildiz, 2024).

Kinesin-14 family members (K-14s) are minus-end directed motors that function in spindle assembly by cross-linking and sliding MTs (She and Yang, 2017), and contribute to pole focusing during both meiosis and mitosis (Endow *et al*., 1990; McDonald and Goldstein, 1990; Hatsumi and Endow, 1992; Endow *et al*., 1994; Walczak *et al*., 1997; Goshima *et al*., 2005; Syrovatkina and Tran, 2015; Higgins *et al*., 2016) and to spindle length control (Saunders *et al*., 1997; Matuliene *et al*., 1999; Mountain *et al*., 1999; Sharp *et al*., 2000; Cai *et al*., 2009; Hepperla *et al*., 2014; Yukawa *et al*., 2018). Spindle length regulation may be attributed in part to Kinesin-14 antagonism with the Kinesin-5 family that separates spindle poles (Saunders *et al*., 1997; Mountain *et al*., 1999; Sharp *et al*., 2000; Yukawa *et al*., 2018). *In vitro,* K-14s can provide an inward sliding force on anti-parallel MTs opposite that of Kinesin-5 outward sliding forces, potentially acting as a brake (Peterman and Scholey, 2009; Hentrich and Surrey, 2010; Reinemann *et al*., 2018). K-14s are also implicated in parallel MT cross-linking and sliding in the spindle (Matuliene *et al*., 1999; Braun *et al*., 2009; Cai *et al*., 2009; Hepperla *et al*., 2014), which may help organize MTs and assist Kinesin-5 outward sliding forces (Henkin *et al*., 2022). Kinesin-14 function is also essential for clustering centrosomes to prevent multipolar divisions and subsequent cell death in cells with centrosome amplification (Kwon *et al*., 2008; Ganem *et al*., 2009) or endoreplication (Chen *et al*., 2016) due to severe aneuploidy, and is proposed to be the motorized force to pull centrosomes together (Rhys *et al*., 2018).

K-14s cross-link both anti-parallel and parallel MTs, which requires both the motor and tail domains (Braun *et al*., 2009; Fink *et al*., 2009; Hentrich and Surrey, 2010; Braun *et al*., 2017) wherein tail binding to MTs is independent of the motor domain (Chandra *et al*., 1993; Hentrich and Surrey, 2010; Braun *et al*., 2017). The tail of *Drosophila* Kinesin-14 Ncd contains two MT binding domains with different MT affinities (Karabay and Walker, 1999; Szczęsna and Kasprzak, 2016), which may make multiple contacts on tubulin dimer (Wendt *et al*., 2003) and appears to rely in part on the acidic C-terminal tails of tubulin (Karabay and Walker, 2003; Furuta and Toyoshima, 2008). *In vitro*, K-14s robustly slide anti-parallel MTs but slide parallel MTs less well or statically cross-link them (Fink *et al*., 2009; Hentrich and Surrey, 2010). MTs in an anti-parallel orientation are slid in the same direction because MT sliding forces would be in the same direction regardless of the motor orientation (Fink *et al*., 2009), suggesting there could be coordination between the motors for anti-parallel MT sliding. However, studies suggest that the density of the motors (Hentrich and Surrey, 2010; Braun *et al*., 2017) or diffusibility of the tail (Lüdecke *et al*., 2018) are inversely correlated with anti-parallel sliding velocity. Little is known about how K-14s statically cross-link parallel MTs, but it is likely due to antagonzing sliding activities of randomly oriented motors within parallel cross-links that exert forces in opposite directions resulting in balanced forces that impede robust sliding (Fink *et al*., 2009; Hentrich and Surrey, 2010).

Given the necessity of the tail domain for MT cross-linking and sliding in the Kinesin-14 family and their roles in spindle assembly and bipolar division, it is important to understand how they differentially cross-link and slide both anti-parallel and parallel MTs *in vitro,* and how these sliding abilities contribute to their physiological functions. If the presence of two MT binding domains in the Kinesin-14 tail is a conserved feature, as suggested by *Drosophila* Ncd (Karabay and Walker, 1999; Szczęsna and Kasprzak, 2016), this could imply that each MT binding domain regulates one type of MT orientation or that their MT affinities might regulate cross-linking and sliding. To address these questions, we identified two MT binding domains in the tail of *Xenopus* Kinesin-14 XCTK2 and systematically characterized their MT affinities and their MT cross-linking and sliding abilities to correlate these activities to spindle assembly and morphology. We found that the two MT binding domains have different MT affinities with the weaker MBD1 imparting increased anti-parallel sliding velocity that correlated with shorter spindles, whereas the tighter MBD2 robustly cross-linked parallel MTs that was important for robust spindle localization and spindle assembly. These findings provide a mechanism by which the affinity of each MT binding domain balances distinct aspects of Kinesin-14 cross-linking and sliding, which can be correlated to their various biological functions.

## Results

### The XCTK2 tail contains two microtubule binding domains, MBD1 and MBD2

Our previous studies showed that importin α/β, effectors of the RanGTP gradient around chromosomes, preferentially inhibit *Xenopus* XCTK2 anti-parallel MT cross-linking and overall MT sliding (Ems-McClung *et al*., 2020), suggesting biochemical differences may exist between the mechanisms of anti-parallel versus parallel MT crosslinking and sliding. In addition, given the existence of two MT binding domains in the tail of *Drosophila* Ncd, one possibility is that there are two separate MT binding domains in the tail of K-14s that recognize two distinct MT orientations and/or regulate MT sliding in which one MT binding domain is regulated by importin α/β. To investigate this idea, we generated a deletion series from the N-terminus (N1-N3) and the C-terminus (C1-C3) of the XCTK2 tail domain (Figure 1, A and B), purified the proteins (Figure S1A), and tested them in MT co-sedimentation assays in the absence and presence of importin α/β (Figure S2A). MT binding relative to the full-length YPet-Tail was reduced for YPet-Tail N1 (p<0.05) but remained equivalent for YPet-Tail N2 (p=0.999) and YPet-Tail N3 (p=0.152) (Figure 1C). Of the C-terminal constructs, all three had MT binding that was indistinguishable from the full-length tail (YPet-Tail C1, p=0.531; YPet-Tail C2, p=0.249; and YPet-Tail C3, p>0.999). Importin α/β significantly inhibited MT binding of the full-length YPet-Tail (p<0.05), YPet-Tail N1 (p<0.0001), YPet-Tail N2 (p<0.0001), and YPet-Tail N3 (p<0.0001), which all contain the NLS of XCTK2 (Figure 1, B and C). In contrast, MT binding of the YPet-Tail C1 was not significantly inhibited by importin α/β, whereas with YPet-Tail C2 and YPet-Tail C3 MT binding was slightly inhibited (p<0.01). We suspect that the reduction in MT binding for YPet-Tail C2 and C3 is due to the predicted simple NLS in these proteins, which is not a functional NLS (Ems-McClung *et al*., 2004; Cai *et al*., 2009). Consistent with this idea, importin α/β affinity assays using Förster resonance energy transfer (Ems-McClung and Walczak, 2020) with importin α-CyPet/importin β and YPet-Tail, YPet-Tail N3, YPet-Tail C2, or YPet-Tail C3 (Figure S2B) show that YPet-Tail N3 has a lower apparent *K*_d_ and lower *B*_max_ compared to YPet-Tail, p<0.05, suggesting a slightly higher importin α/β affinity, but YPet-Tail C2 and YPet-Tail C3 have about an eight-fold higher apparent *K*_d_ and about six-fold lower *B*_max_ compared to YPet-Tail, p<0.0001 (Figure 1D), consistent with an inability to robustly bind importin α/β. These results show that there are two separable MT binding domains in the tail of XCTK2, one of which is regulated by importin α/β through the NLS. Because YPet-Tail N3 and YPet-Tail C2 fragments had the most robust and reproducible MT binding, we used these for further analysis.

**Figure 1.**
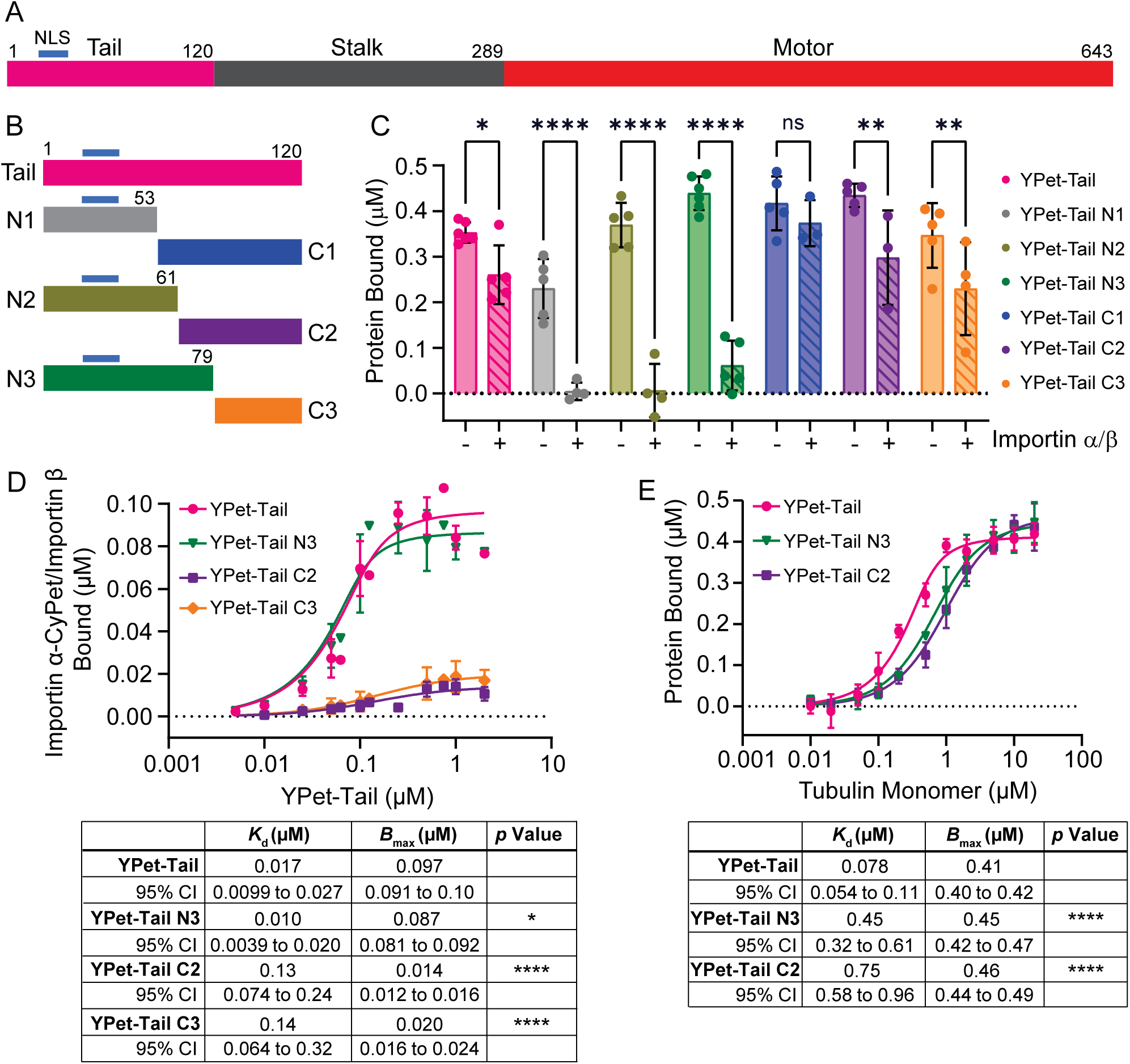
(A) Schematic of domain organization and amino acid numbers of the Kinesin-14 XCTK2 drawn to scale. The tail (pink) is at the N-terminus, the stalk (gray) is in the middle, and the motor (red) is at the C-terminus. The NLS is indicated by a blue bar. (B) Schematic of deletion series of the tail drawn to scale with the NLS indicated by a blue bar. (C) Bar graph of the mean ± SD of 0.5 μM YPet-Tail fragments bound to 5 μM MTs in the absence (solid) or presence (hashed) of 2 μM importin α/β in a MT co-sedimentation experiment and analyzed by a two-way ANOVA. N=3-6 independent experiments. (D) Importin α/β affinity assays using Förster resonance energy transfer with 0.1 μM importin α-CyPet/0.4 μM importin β and 0-2 μM YPet-Tail, YPet-Tail N3, YPet-Tail C2, or YPet-Tail C3 fit with a quadratic equation for binding and each concentration graphed as the mean ± SD from 4-7 independent experiments. (E) MT affinity curves of 0.5 μM YPet-Tail, N3 or C2 graphed as the mean ± SD of protein bound to increasing concentrations of MTs reported as tubulin monomer (0-20 μM) and fit to the quadratic equation for saturable binding. N=3-8 independent experiments. The apparent affinity curves between YPet-Tail and N3, C2, and/or C3 proteins were compared with an extra sum-of-squares F-test. ns, not significant; *, p<0.05; **, p<0.01; ****, p<0.0001.

If the MT binding sites within the XCTK2 tail function as separable binding sites, then we would predict that each MT binding domain could bind to a site on tubulin dimer within the MT polymer such that two MT binding sites would exist per tubulin dimer. To determine the stoichiometry of MT binding, we measured the MT binding of YPet-Tail, YPet-Tail N3, and YPet-Tail C2 using MT co-sedimentation with increasing concentrations of MTs (Figure S2C) and fit the data to the stoichiometric quadratic equation (Bell *et al*., 2017). The stoichiometry model indicated that at least two full-length YPet-Tail proteins bound per tubulin dimer because the stoichiometry coefficient, *s*, was 2.4 (Figure S2D). In contrast, the stoichiometry model indicated that YPet-Tail N3 and YPet-Tail C2 bound once per tubulin dimer, *s* = 1.1. One possibility is that each MT binding domain binds to a tubulin monomer. If this is true, then the MT binding of the tail proteins should fit the quadratic equation plotted per tubulin monomer (Figure 1E). The MT binding of the YPet-Tail, YPet-Tail N3, and YPet-Tail C2 proteins fit this model well with YPet-Tail having a six-fold higher apparent MT affinity compared to YPet-Tail N3, p<0.0001, and having eight-fold higher apparent MT affinity than YPet-Tail C2, p<0.0001 (Figure 1E). Furthermore, the apparent affinity for MT binding of YPet-Tail N3 was about two-fold higher than YPet-Tail C2 (p<0.05), suggesting YPet-Tail N3 might have a slightly higher MT affinity. Together, these results suggest that the tail of XCTK2 contains an N-terminal MT binding domain that can be inhibited by importin α/β (MBD1) and a C-terminal MT binding domain that is not inhibited by importin α/β (MBD2), both of which bind a tubulin monomer within the MT.

### MBD2 has higher MT affinity than MBD1 and is insensitive to importin α/β

To identify the key residues important in MT binding we mutated lysine and arginine residues (K/R) to alanine in the N3 and C2 constructs (Figures 2A and S1B) and assayed them for their ability to bind MTs (Figure 2, B and C). Our previous studies indicated that mutation of the NLS reduced MT binding in the full-length tail (Ems-McClung *et al*., 2004); thus we compared MT binding of the NLSb partial mutation to the full mutation of both NLSa and NLSb (MBD1^mut^) in YPet-Tail N3 (Figure 2, A and B). YPet-Tail N3-MBD1^mut^ containing six mutated K/R residues had a 10-fold higher *K*_d_ and an eight-fold lower *B*_max_, indicating a drastically reduced apparent MT affinity compared to YPet-Tail N3, p<0.0001 (Figure 2B). To inhibit the MT binding of MBD2, we systematically mutated three different sets of K/R residues within CyPet-Tail C2 (Figure 2, A and C). Ultimately, mutation of eight K/R residues in CyPet-Tail C2-MBD2^mut^ increased the *K*_d_ four-fold and reduced the *B*_max_ four-fold, significantly inhibiting the apparent MT affinity relative to CyPet-Tail C2, p<0.0001 (Figure 2C).

**Figure 2.**
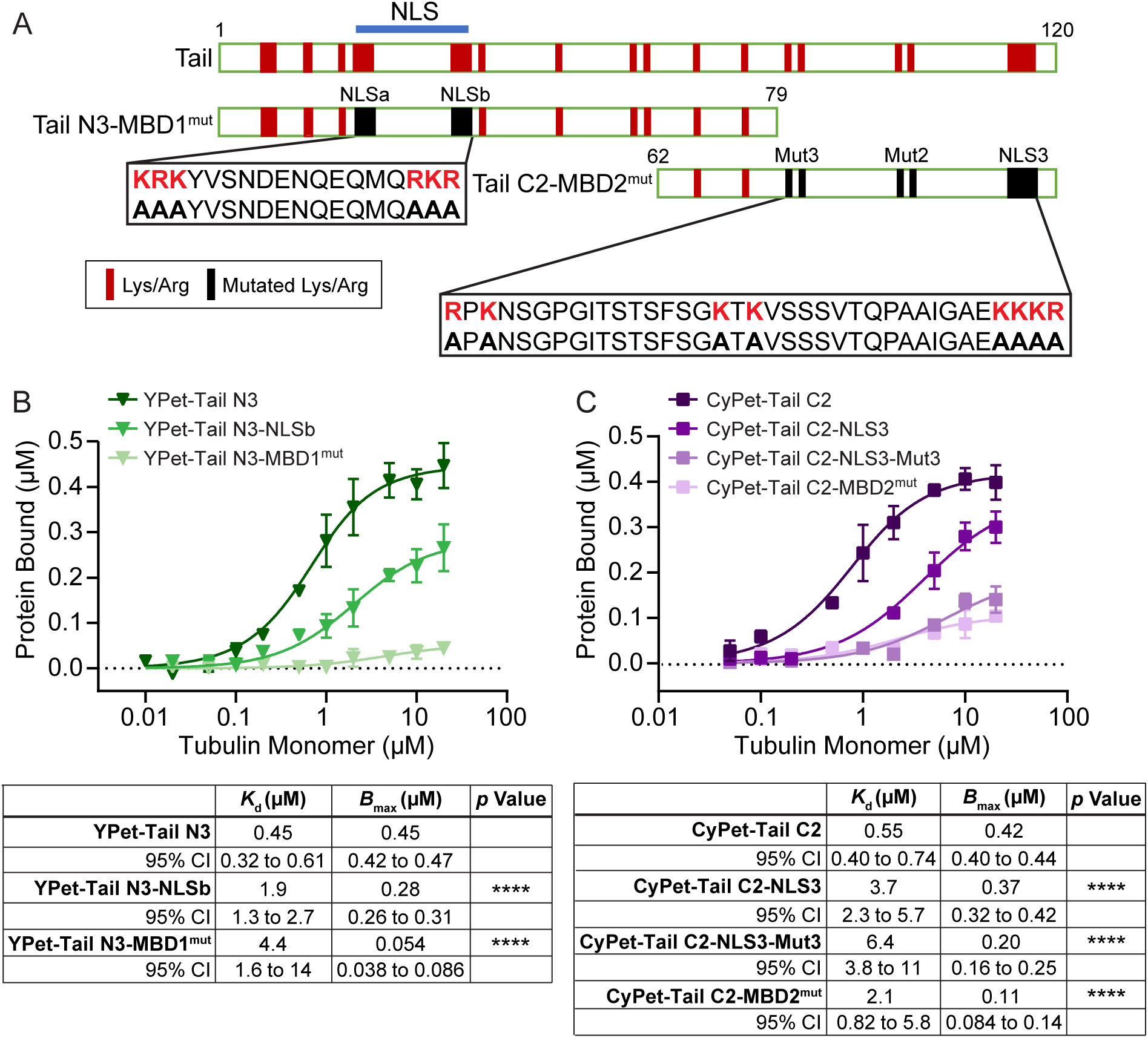
(A) Schematic of the XCTK2 Tail protein and mutated Tail N3 or Tail C2 protein amino acid sequences drawn to scale with K/R indicated in red and mutated K/R in black. The NLS is indicated with a blue bar. MT affinity curves from MT sedimentation assays comprising 0.5 μM wild-type or mutated YPet-Tail N3 (B) or CyPet-Tail C2 (C) graphed as the mean ± SD of tail protein bound to increasing concentrations of MTs reported as tubulin monomer (0-20 μM) and fit to the quadratic equation for saturable binding. The apparent affinity curves between wild-type YPet-Tail N3 or CyPet-Tail C2 proteins and each mutant were compared with an extra sum-of-squares F-test. N=3-6 independent experiments. ****, p<0.0001.

To address the relative MT affinities of the two MBDs in the context of the full-length XCTK2 tail, we incorporated the MBD1^mut^ and MBD2^mut^ mutations individually and together into the full tail domain (Figures 3A and S1C) and assessed their apparent MT binding affinities (Figure 3B). MT binding of YPet-Tail-MBD1^mut^ had a six-fold lower apparent affinity than wild-type YPet-Tail, p<0.0001, and YPet-Tail-MBD2^mut^ had a 27-fold lower apparent affinity, p<0.0001. Congruent with mutating both MT binding domains, YPet-Tail-MBD1-2^mut^ did not effectively bind MTs and had an apparent affinity 423-fold lower than YPet-Tail, p<0.0001. These results suggest that MBD2 has a higher MT affinity than MBD1 in the context of the full-length tail since YPet-Tail-MBD1^mut^ with functional MBD2 had a four-fold higher apparent affinity than YPet-Tail-MBD2^mut^, p<0.0001 (Figure 3B). These results indicate that we have identified the key residues that mediate MT binding of each MT binding domain within the XCTK2 tail.

**Figure 3.**
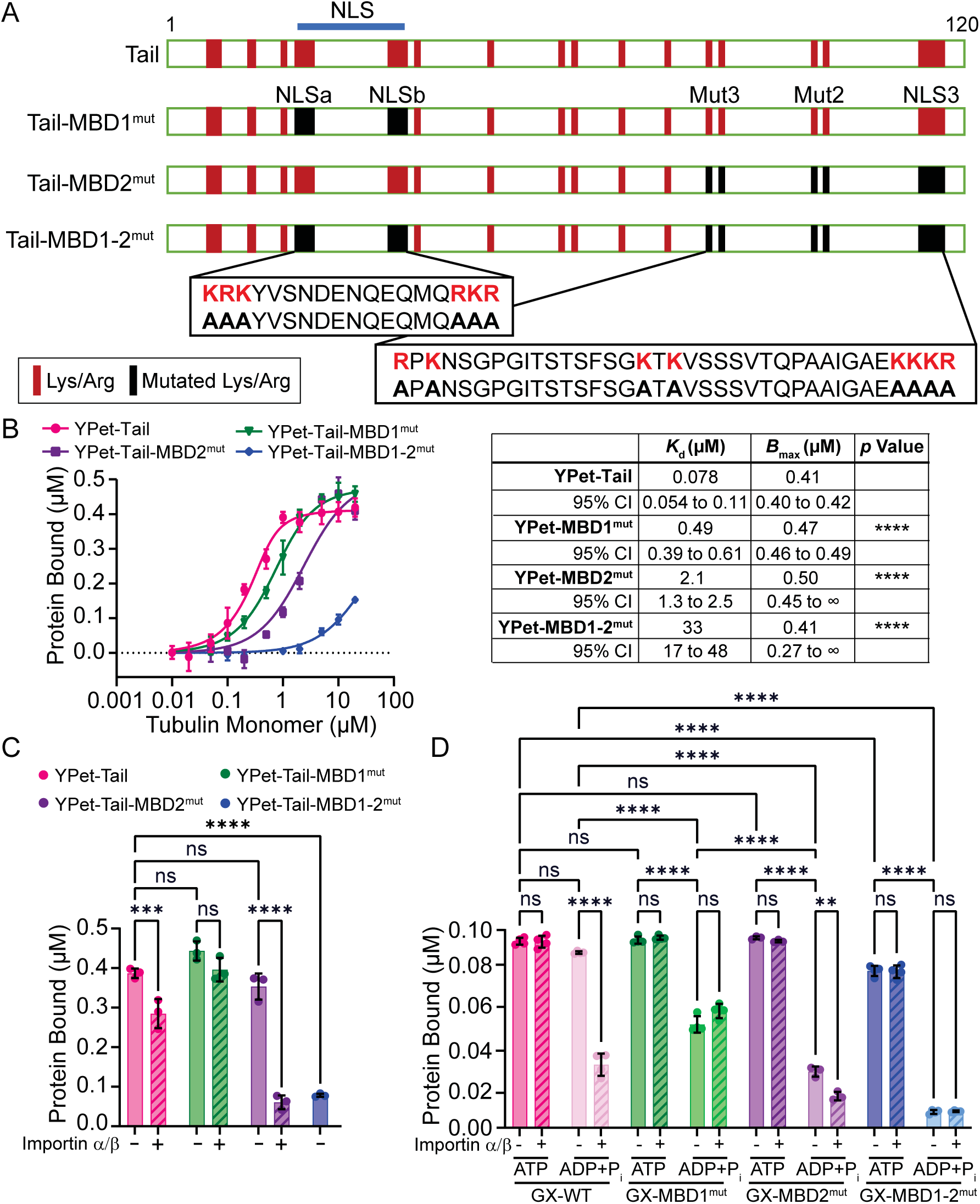
(A) Schematic of the wild-type and mutant XCTK2 Tail protein amino acid sequences drawn to scale with K/R residues indicated in red and mutated K/R in black. The NLS is indicated with a blue bar. (B) MT affinity curves from MT co-sedimentation assays of 0.5 μM wild-type or MBD mutated full-length tail proteins graphed as the mean ± SD of tail protein bound to increasing concentrations of MTs reported as tubulin monomer (0-20 μM) and fit to the quadratic equation for saturable binding. The apparent affinities between YPet-Tail and each MBD mutant were compared with an extra sum-of-squares F-test. N=3-8 independent experiments. (C) Bar graph of the mean ± SD amount of 0.5 μM full-length YPet-Tail or MBD mutant proteins bound to 5 μM MTs in the absence (solid) or presence (hashed) of 2 μM importin α/β in a MT co-sedimentation experiment and compared using a two-way ANOVA. N=3 independent experiments. (D) Bar graph of the mean ± SD amount of 0.1 μM full-length GFP-XCTK2 wild-type or MBD mutant proteins in the absence (solid) or presence (hashed) of 0.4 μM importin α/β bound to 1 μM MTs in 2 mM MgATP (dark) or 2 mM MgADP+20 mM P_i_ (light) in a MT co-sedimentation experiment and compared using a three-way ANOVA. N=3-4 independent experiments. ns, not significant; **, p<0.01; ***, p<0.001; ****, p<0.0001.

XCTK2 tail MT binding is regulated by importin α/β (Ems-McClung *et al*., 2004), and this regulation is important physiologically because XCTK2 association with the spindle is spatially regulated by importin α/β through the RanGTP gradient (Weaver *et al*., 2015; Ems-McClung *et al*., 2020). To test how importin α/β affect MT binding of the full-length tail with the MBD mutations, YPet-Tail, YPet-Tail-MBD1^mut^, YPet-Tail-MBD2^mut^, or YPet-Tail-MBD1-2^mut^ were incubated with MTs in the absence or presence of importin α/β in the MT co-sedimentation assay (Figure 3C). Consistent with our initial assay (Figure 1C) and with our previous findings (Ems-McClung *et al*., 2004), YPet-Tail bound MTs well in the absence of importins, but MT binding was reduced in the presence of importin α/β, p<0.001. YPet-Tail-MBD1^mut^ also bound MTs well, but MT binding was insensitive to importin α/β as expected because the K/R residues mutated in MBD1^mut^ constitute the NLS. In contrast, the MT binding of YPet-Tail-MBD2^mut^ was dramatically inhibited by the importins, p<0.0001. Mutation of both MBDs in YPet-Tail-MBD1-2^mut^ resulted in the inability of the tail to bind MTs, p<0.0001, as expected based on its drastically reduced MT affinity (Figure 3B). These results are consistent with MBD1 being the Ran-regulated MT binding domain and that in the presence of importins any MT binding of the tail likely occurs through MBD2.

To examine how the MBD mutations affect MT binding of the dimeric full-length protein, we introduced the MBD1^mut^ and MBD2^mut^ mutations individually and together (MBD1-2^mut^) into full-length GFP-XCTK2, expressed and purified them (Figure S1C), and then performed a MT co-sedimentation assay in 2 mM MgATP or 2 mM MgADP + 20 mM P_i_ in the absence or presence of importin α/β (Figure 3D). In the absence of importins and the presence of MgATP, GFP-XCTK2 (GX-WT), GFP-XCTK2-MBD1^mut^ (GX-MBD1^mut^) and GFP-XCTK2-MBD2^mut^ (GX-MBD2^mut^) bound MTs similarly and well, consistent with the motor domains readily binding MTs in MgATP (Figure 3D, - Importin α/β, ATP). GFP-XCTK2-MBD1-2^mut^ (GX-MBD1-2^mut^), on the other hand, bound MTs 16% less well than GX-WT, p<0.0001. The reduced MT binding of GX-MBD1-2^mut^ indicates that tail MT binding does contribute to the overall protein binding in MgATP, which is consistent with the idea that the tail domain tethers K-14s between cross-linked MTs during their catalytic cycle (Braun *et al*., 2017). To assess the contribution of the tail MT binding domains to MT binding in the absence of motor domain MT binding, we assayed MT binding in the presence of MgADP+P_i_ where the motor domains are predominantly not bound to MTs (Rosenfeld *et al*., 1996; Foster *et al*., 1998). For GX-WT, most of the protein was still bound to MTs in MgADP+P_i_, whereas mutation of either MT binding domain reduced MT binding ≥50% compared to its MT binding in MgATP, p<0.0001, and compared to GX-WT in MgADP+P_i_, p<0.0001 (Figure 3D, - Importin α/β). These results suggest that mutation of either one of the tail MT binding domains hinders the ability of the tail to tether the protein to MTs during its catalytic cycle *in vitro*. In addition, MT binding of GX-MBD1^mut^ in MgADP+P_i_ was approximately two-fold higher than GX-MBD2^mut^, p<0.0001, consistent with the MT affinities determined above that indicated functional MBD2 has tighter MT affinity than functional MBD1 (Figure 3, B and D). MT binding of GX-MBD1-2^mut^ was minimal in MgADP+P_i_, p<0.0001 (Figure 3D), consistent with our finding that these mutations abolish MT binding by the tail (Figure 3, B and C).

To assess how the importins regulate MT binding of full-length GX-WT and the MT binding domain mutants during their catalytic cycle, we added the importins in excess to full-length protein in the MT co-sedimentation assay in the presence of MgATP or MgADP+P_i_ (Figure 3D, + Importin α/β). Addition of the importins had no effect on the MT binding of any of the proteins in MgATP compared to without importins, consistent with MT binding occurring predominantly through the motor domain in this nucleotide condition. However, the presence of the importins reduced GX-WT MT binding 64% in the presence of MgADP+P_i_ compared to without importins, p<0.0001, consistent with MBD1 MT binding being inhibited. Importin addition to GX-MBD1^mut^ or GX-MBD1-2^mut^ in MgADP+P_i_ did not further inhibit MT binding due to the mutated NLS in MBD1. GX-MBD2^mut^ MT binding in the presence of MgADP+P_i_ and importins was reduced by 43% compared to without importins, p<0.01, consistent with importin mediated inhibition of MBD1 MT binding. Together our results support the idea that MT binding by the tail domain is important to tether the full-length protein to MTs during its catalytic cycle, that the two tail MT binding domains have different MT affinities (MBD2 > MBD1), and that MT binding by MBD1 is regulated by the Ran pathway.

### MBD1 and MBD2 differentially promote spindle assembly and morphogenesis

To understand how the differences in MT affinity of the two different MT binding domains impact XCTK2 physiological function, we examined how full-length GX-WT or the MBD mutant proteins affected spindle assembly in *Xenopus* egg extracts (Figure 4A). With the addition of a 2.5-fold molar excess of protein, both GX-WT and GX-MBD1^mut^ localized heavily to spindles assembled in CSF extracts (Figures 4A and S3A) and stimulated the formation of bipolar spindles relative to spindle intermediates (asters and half spindles) that was approximately two-fold over that of GFP control, p<0.01 and p<0.05 respectively (Figure 4B), consistent with our previous findings that over addition of XCTK2 increases the efficiency of spindle assembly (Walczak *et al*., 1997; Cai *et al*., 2009; Weaver *et al*., 2015). GX-MBD2^mut^ increased spindle assembly by 52% over GFP control and 27% less than GX-WT but was not significantly different from either GFP control or GX-WT (Figure 4B), which suggests a partial stimulation of spindle assembly. GX-MBD2^mut^ did not localize well to spindle assembly structures (Figure 4A), which may account for its less efficient stimulation of spindle assembly. GX-MBD1-2^mut^ was unable to stimulate spindle assembly over GFP control (Figure 4B) and localized poorly to spindle assembly structures (Figure 4A), consistent with our previous results with GFP-XCTK2 (122-643) lacking the tail domain (Weaver *et al*., 2015). These results suggest that MBD1 is dispensable for the stimulation of spindle assembly and that the tighter MT affinity of MBD2 is sufficient for the full-length protein to bind to spindle MTs and stimulate spindle assembly in extracts.

**Figure 4.**
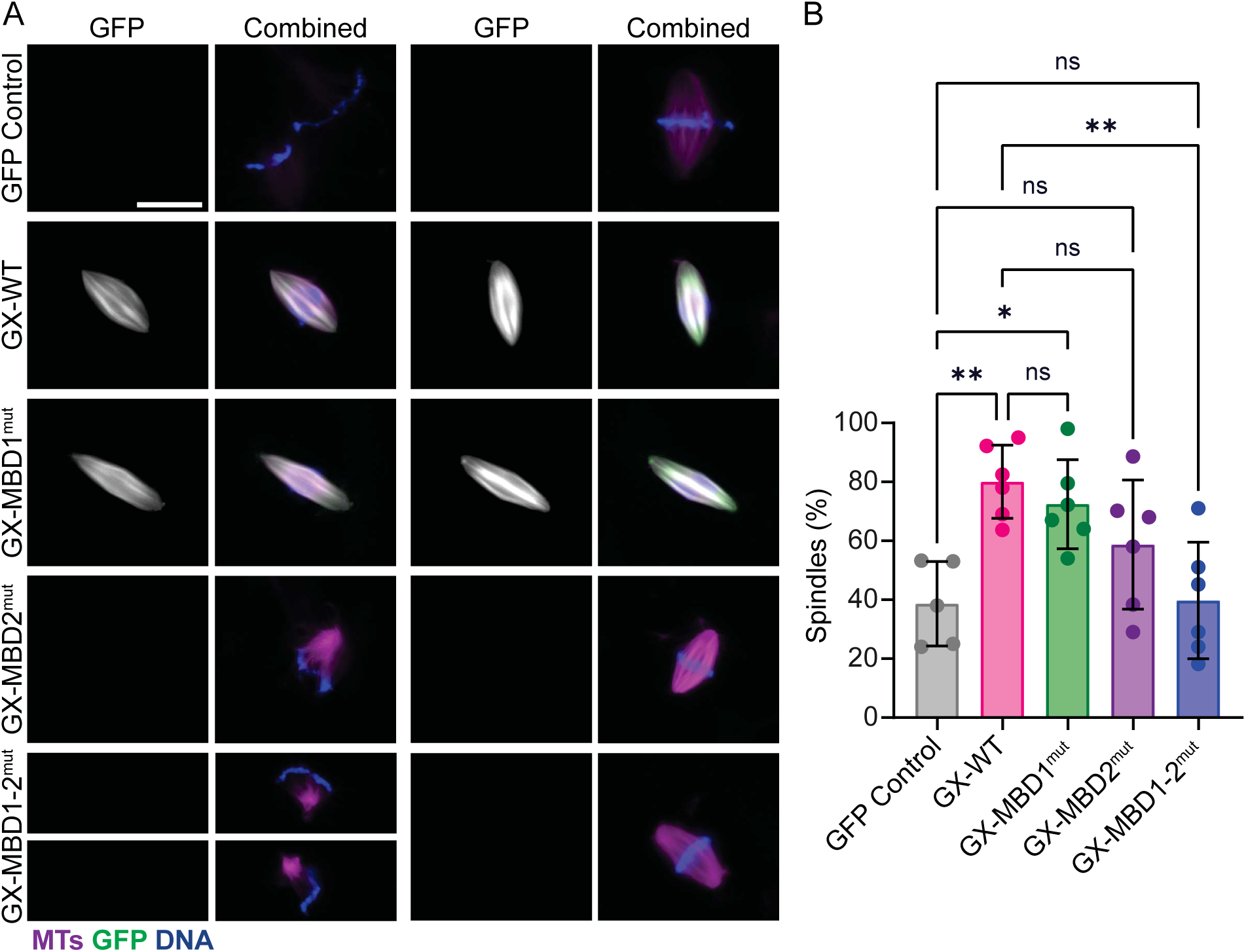
(A) Representative images of spindle structures assembled in CSF extracts in which 50 nM GFP, GX-WT, GX-MBD1^mut^, GX-MBD2^mut^, or GX-MBD1-2^mut^ were added. The individual GFP channel is grayscale. For the combined n images, MTs are magenta, GFP protein is green, and DNA is blue. Scale bar, 20 μm. (B) Percentages of bipolar spindles formed by the over addition of GX proteins as described in (A). Data is from 5-6 independent CSF extracts graphed as the mean ± SD. Spindle numbers: GFP control n=615, GX-WT n=603, GX-MBD1^mut^ n=599, GX-MBD2^mut^ n=609, GX-MBD1-2^mut^ n=640. ns, not significant; *, p<0.05; **, p<0.01.

To investigate how differences in tail MT binding impacts protein localization and spindle morphology, we assembled cycled spindles, which are more uniform and robust, in the presence of a 2.5-fold molar excess of GX-WT, GX-MBD1^mut^, or GX-MBD2^mut^ (Figures 5A and S3B). We did not assess GX-MBD1-2^mut^ since it did not stimulate CSF spindle assembly, and because it localized poorly to spindles. Spindles assembled in the presence of GX-WT appeared to have more MTs compared to GFP control (Figure 5A, first row), suggesting increased MT polymer. In contrast, spindle MTs with GX-MBD1^mut^ and GX-MBD2^mut^ did not appear as bright as GX-WT spindle MTs (Figure 5A). To determine whether the over addition of the GX proteins increased spindle MT polymer, we measured the total integrated MT intensities of the spindles in each condition (Figure 5B). GX-WT spindles had higher MT intensities than the GFP control or the MBD mutants, p<0.0001, suggesting that GX-WT over addition increased total MT polymer. Over addition of the MBD mutants did not increase total MT intensities significantly, suggesting that they do not elevate overall MT polymer levels. Visually, GX-WT and GX-MBD1^mut^ localized well to spindles, whereas GX-MBD2^mut^ localized less effectively (Figure 5A, second row), consistent with the observations in the CSF spindle assembly reactions. To compare the localization of the added proteins, we used the integrated GFP intensities because the spindle areas with the over addition of the GX proteins were reduced compared to GFP control (Figure S3C). GX-MBD1^mut^ localization on spindles was 45% lower than GX-WT (p<0.01) (Figure 5C), which aligns with previous findings that RanGTP recruits endogenous XCTK2 to the spindle via the NLS, which resides within MBD1 (Ems-McClung *et al*., 2020). GX-MBD2^mut^ accumulated 76% less than GX-WT (p<0.0001) and 57% less than GX-MBD1^mut^, p<0.0001, suggesting that the higher MT affinity of functional MBD2 in GX-WT and GX-MBD1^mut^ may help tether the protein to the MTs of the spindle or that other mechanisms govern spindle localization via the MT binding domains.

**Figure 5.**
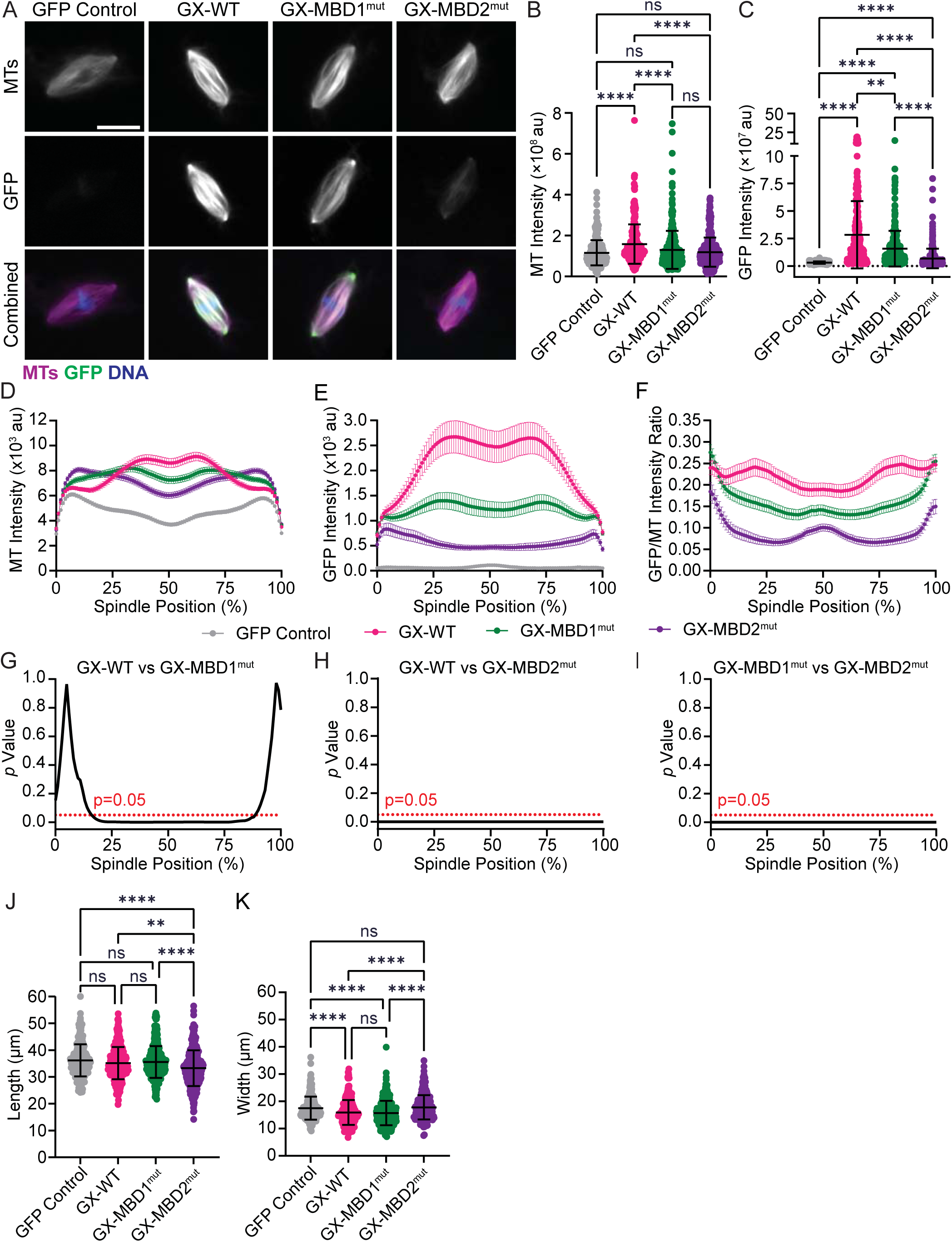
(A) Representative images of spindles assembled in cycled extracts in which 50 nM GFP, GX-WT, GX-MBD1^mut^, or GX-MBD2^mut^ were added. Individual MT and GFP channels are grayscale. For the combined images, MTs are magenta, GFP protein is green, and DNA is blue. Scale bar, 20 μm. Measured integrated MT intensity (B) and integrated GFP intensity (C) plotted with the mean ± SD from N=3 independent experiments. (D-F) Line scans normalized to spindle length of integrated MT intensity (D), integrated GFP intensity (E), or the ratio of integrated GFP to MT intensity (F), plotted as the mean ± SEM. (G-I) *p* value at each spindle position between GX-WT and GX-MBD1^mut^ (G) or GX-MBD2^mut^ (H), and between GX-MBD1^mut^ and GX-MBD2^mut^(I). Spindle length (J), and spindle width (K) are plotted with the mean ± SD. Spindle numbers: GFP control n=273-274, GX-WT n=296, GX-MBD1^mut^ n=305-308, GX-MBD2^mut^ n=300-301. ns, not significant; **, p<0.01; ****, p<0.0001.

To ask whether the mutations affected the distribution of the proteins on the spindle, we did a line scan analysis from pole-to-pole to assess the distribution of MT and GFP intensities across the spindle (Figure 5, D-F). Spindles with GFP control protein showed increased MT intensity near the poles relative to the body of the spindle. Spindles with GX-WT had increased MT polymer levels over that of GFP control, especially in the body of the spindle (Figure 5D). GX-MBD1^mut^ addition also increased the MT polymer levels in the body of the spindle over that of GFP control, but MT intensities were more evenly distributed across the spindle. The overall MT intensities of spindles with GX-MBD2^mut^ were also higher than spindles with GFP control but were lower than GX-WT and GX-MBD1^mut^ in the body of the spindle. The differences in the levels of GFP intensity of spindles with GX-WT, GX-MBD1^mut^, and GX-MBD2^mut^ correlated with the differences in intensities of MTs in the spindles (Figure 5E). To ask whether the increased amounts of the added GX proteins on the spindle were due to the changes in MT polymer and distribution, we calculated the ratio of the integrated GFP intensities to integrated MT intensities along each spindle (Figure 5F). The ratio of GX-WT to MT intensity was lower in the middle of the spindle and higher near the poles, which is consistent with the endogenous localization of XCTK2 described previously (Walczak *et al*., 1997). GX-MBD1^mut^ and GX-MBD2^mut^ were more evenly distributed across the spindle at levels less than GX-WT. To compare the difference in the distribution of the proteins across the spindle, we performed statistical tests on the GFP/MT intensity ratios for each pair of proteins at each position along the spindle and plotted the *p* values (Figure 5, G-I). GX-WT was higher throughout the spindle compared to GX-MBD1^mut^ except at the poles, whereas GX-WT was higher than GX-MBD2^mut^ across the entire spindle (Figure 5, G and H). In addition, GX-MBD1^mut^ was higher than GX-MBD2^mut^ across the entire spindle (Figure 5I). These results suggest that both MBDs are needed for the increased GX-WT accumulation on the spindle, and that the levels of GX protein localization correlate with the affinity of the tail for MTs.

When examining morphological differences of the spindles, we noted that spindles with GX-WT and GX-MBD1^mut^ appeared narrower than spindles with control GFP control spindles and that spindles with GX-MBD2^mut^ appeared shorter than control spindles (Figure 5A), suggesting that the different MT binding domains contributed differentially to spindle morphogenesis. To characterize the spindle morphologies, we measured the length and width of the spindles (Figure 5, J and K). Spindles with GX-WT and GX-MBD1^mut^ were similar in length to control spindles (Figure 5J) but were approximately 2 μm thinner (p<0.0001) (Figure 5K), suggesting that the over addition of the tighter MT binding of MBD2 may bundle or more stably cross-link MTs within the spindle, resulting in thinner spindles. Consistent with this idea, the spindle areas of spindles with GX-WT and GX-MBD1^mut^ were smaller than GFP control, p<0.0001, but equivalent to one another (Figure S3C). Despite its reduced localization, spindles with GX-MBD2^mut^ were about 3 μm shorter than control spindles (p<0.0001) and about 2 μm shorter than spindles with GX-WT (p<0.01) but were as wide as control spindles (Figure 5, J and K). The reduced length of spindles with GX-MBD2^mut^ correlated with their reduced spindle areas compared to GFP control spindles, p<0.05 (Figure S3C). The shorter spindle lengths of spindles with GX-MBD2^mut^ could be due to better anti-parallel sliding and/or hindered tight MT cross-linking due to the MBD2 mutations. Together, these results suggest the MBDs may regulate spindle length through two modalities: anti-parallel sliding modulated via MBD1 and stable MT cross-linking by MBD2.

### Dual tail MT binding domains control anti-parallel MT sliding velocity (MBD1) and the extent of parallel MT cross-linking (MBD2)

Our previous studies showed that importin α/β preferentially inhibited anti-parallel MT cross-linking and MT sliding (Ems-McClung *et al*., 2020), which may indicate that the two MBDs in the tail modulate different types of cross-links where MBD1 is needed for anti-parallel MT cross-linking, and MBD2 is needed for parallel MT cross-linking. Alternatively, the strength of tail MT binding or the diffusibility of the tail has been suggested to modulate the robustness of MT sliding (Braun *et al*., 2017; Lüdecke *et al*., 2018), which might suggest that the weaker MT affinity of MBD1 may contribute to robust MT sliding, which is more prevalent in anti-parallel cross-links, whereas the tighter MT affinity of MBD2 may contribute to the tighter cross-linking and reduced sliding of parallel MTs.

To examine how the differential MT affinities of the MT binding domains affect MT cross-linking and sliding, we performed timelapse microscopy of polarity marked MTs with GX-WT, GX-MBD1^mut^, GX-MBD2^mut^, or GX-MBD1-2^mut^ undergoing cross-linking and sliding on template MTs (Figure 6, A and B) and scored the number of cargo MTs and their behavior. Because mutation of the tail MT binding domains reduced the overall number of MT cross-links, we used twice the amount of mutant protein relative to WT (10 nM versus 5 nM) to have sufficient cross-links to analyze. With GX-WT, GX-MBD1^mut^, or GX-MBD2^mut^, cargo MTs cross-linked to template MTs and/or glided across the surface of the coverslip, whereas with GX-MBD1-2^mut^, the cargo MTs were either statically cross-linked or stuck to the slide without any directed movement. When GX-WT, GX-MBD1^mut^, or GX-MBD2^mut^ glided on the surface of the chambers, the long extension of the polarity marked MTs always led, consistent with XCTK2 being a minus-end directed motor like other Kinesin-14 proteins (Chandra *et al*., 1993; Stewart *et al*., 1993; Furuta *et al*., 2008; Fink *et al*., 2009; Hentrich and Surrey, 2010; Duan *et al*., 2012; Braun *et al*., 2017). When scoring the number of cargo MTs per field of view (Figure 6B), GX-WT had the most cargo MTs present, whereas GX-MBD1^mut^ and GX-MBD2^mut^ had about 60% fewer cargo MTs, p<0.01, and GX-MBD1-2^mut^ had very few cargo MTs, p<0.0001 (Figure 6C). Because GX-MBD1-2^mut^ had very few cargo MTs, we did not further analyze movies with this protein. These results are consistent with our biochemical characterization of the tail in which mutation of either MT binding domain reduced the MT binding affinity compared to the wild-type tail, and mutation of both MBDs dramatically inhibited the ability of the tail to bind MTs and thus prevented MT cross-linking.

**Figure 6.**
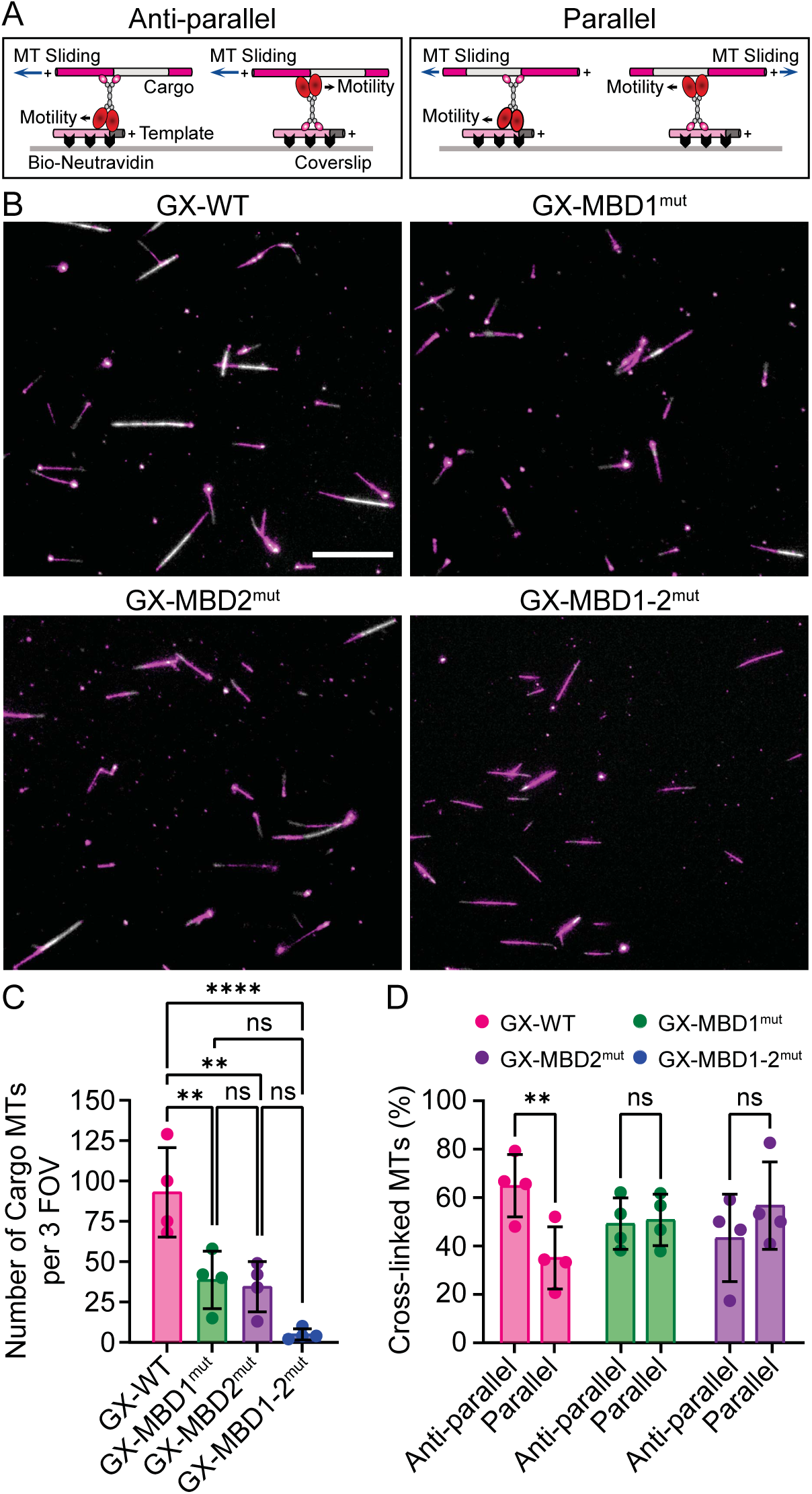
(A) Schematic of the MT cross-linking and sliding assay showing the directionality that the cargo MT can move depending on the orientation and movement of the motor when MTs are cross-linked in an anti-parallel (left) or parallel (right) manner. MT plus-ends are indicated by a plus sign (+), and the direction of motor or cargo MT movement is indicated by an arrow. Polarity marked MT plus ends polymerize faster and either result in a single extension or a longer extension than the minus ends. (B) Representative fields of view of the cross-linking and sliding assays in which 5 nM GX-WT or 10 nM GX-MBD1^mut^, GX-MBD2^mut^, or GX-MBD1-2^mut^ were added. Scale bar, 20 μm. (C) Average number of cargo MTs in three fields of view plotted as the mean ± SD and compared with a one-way ANOVA. GX-WT, n=372; GX-MBD1^mut^, n=155; GX-MBD2^mut^, n=138; GX-MBD1-2^mut^, n=20. (D) Average percentage of cross-linked cargo MTs that were in aligned anti-parallel or parallel MT cross-links plotted as the mean ± SD and compared with a two-way ANOVA. GX-WT, n=107; GX-MBD1^mut^, n=123; GX-MBD2^mut^, n=93. For all data, n is the number of microtubules in N=4 independent experiments. ns, not significant; **, p<0.01; ****, p<0.0001.

To ask how mutation of the tail MT binding domains impacted the orientation and sliding of the MT cross-links, we analyzed similar numbers of cross-linked MTs from four independent experiments for an average of 35 MT cross-links for GX-WT (n=138 total cross-links), 38 MT cross-links for GX-MBD1^mut^ (n=152 total cross-links), and 38 MT cross-links for GX-MBD2^mut^ (n=151 total cross-links). From these MT cross-links, we determined the percentage of MT cross-links that were aligned anti-parallel or parallel (Figure 6D). GX-WT preferentially cross-linked anti-parallel MTs compared to parallel MTs, p<0.01, whereas GX-MBD1^mut^ and GX-MBD2^mut^ crosslinked anti-parallel and parallel MTs in similar proportions (Figure 6D), consistent with previous studies (Fink *et al*., 2009; Hentrich and Surrey, 2010; Ems-McClung *et al*., 2020). The observation that mutation of either MBD does not specify a preference for one type of cross-link goes against the idea that each MT binding domain is responsible for one type of MT cross-link orientation and suggests that the tail does not explicitly determine the orientation of the cross-link.

If the tail MT binding domains do not determine the orientation of MT cross-links, then perhaps they control MT sliding behavior such as the extent or velocity of sliding. Cargo MTs aligned in anti-parallel or parallel MT orientations by GX-WT, GX-MBD1^mut^, or GX-MBD2^mut^ were observed to either slide or be statically cross-linked (Figure 7A and Video S1-3). All three proteins slid the majority of the anti-parallel MT cross-links (75-95%) with the cargo MT plus ends always moving toward the minus ends of the template MTs (Figure 7, A and B and Videos S1-3), consistent with previous work (Fink *et al*., 2009; Hentrich and Surrey, 2010; Braun *et al*., 2017). For aligned parallel MT cross-links that slid, the cargo MTs either slid their minus ends toward the minus ends of the template MTs or slid their plus ends toward the plus ends of the template MTs (Figures 6A and 7A). In the presence of GX-WT and GX-MBD1^mut^, only a small percentage of parallel cargo MTs slid (9-21%) with the remaining being statically cross-linked (Figure 7B), consistent with previous results that most parallel MT cross-links are static (Fink *et al*., 2009; Hentrich and Surrey, 2010; Braun *et al*., 2017). In contrast, GX-MBD2^mut^ slid 51% of the parallel MT cross-links, suggesting that the weakened tail MT affinity modulates the extent of parallel MT sliding (Figure 7B).

**Figure 7.**
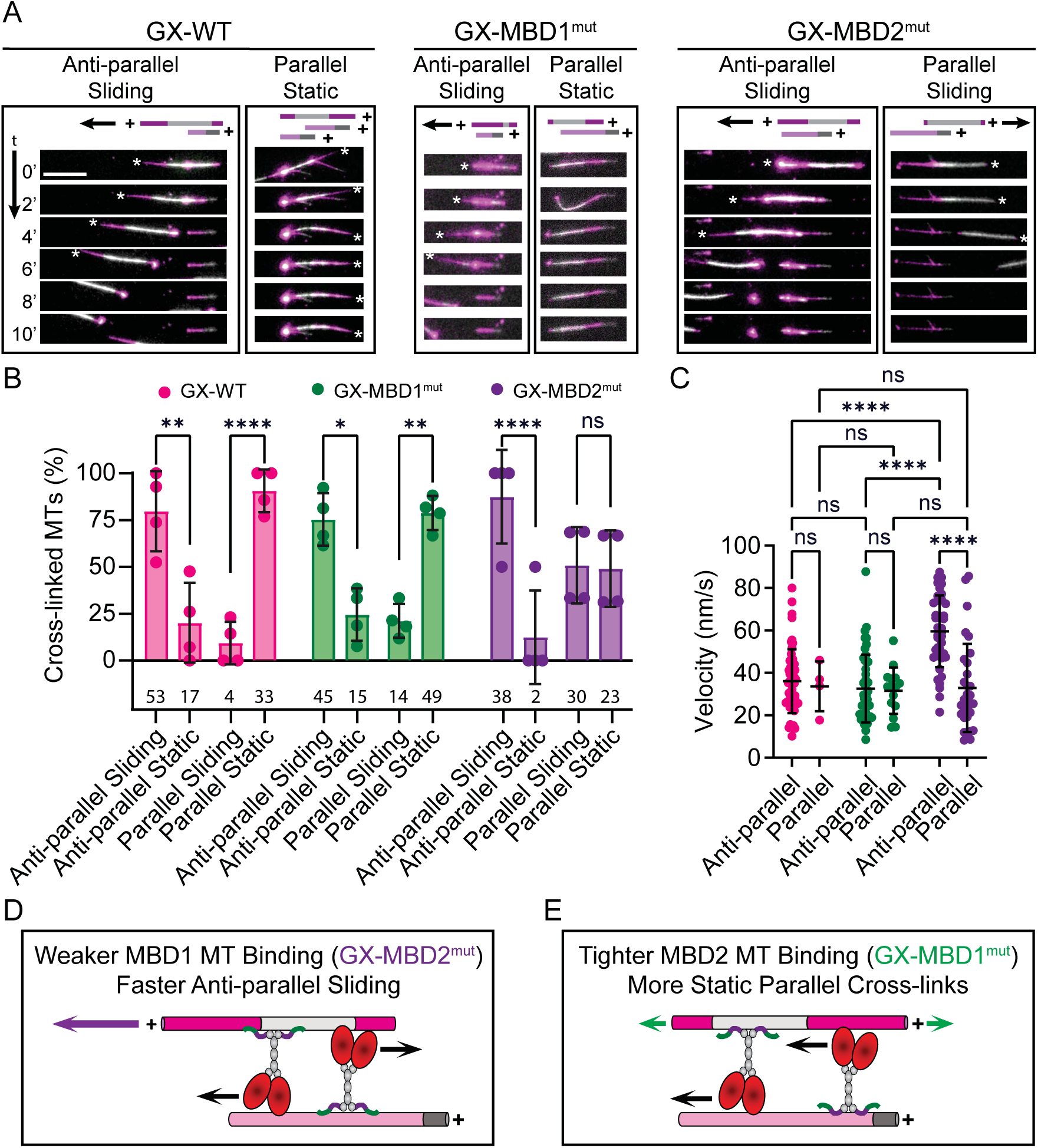
(A) Representative image series at two-minute intervals of aligned anti-parallel and parallel MT cross-linking and sliding events. The schematic above each panel represents the MTs in the corresponding image with the MT plus-ends (+) and the direction of sliding (arrow) indicated. The asterisks in the images indicate the plus end of the long extension of the polarity marked MT over time. Note that in the GX-WT, GX-MBD1^mut^, and GX-MBD2^mut^ anti-parallel and GX-MBD2^mut^ parallel sliding events that the cargo MT slides off the template MT and continues moving by gliding at a faster velocity. Scale bar, 10 μm. (B) Average percentage of aligned cargo MTs that slid or were statically cross-linked in anti-parallel or parallel configurations plotted as the mean ± SD and compared by a two-way ANOVA. Total cargo MTs for each category are indicated in the graph. (C) Average MT sliding velocities of cargo MTs in aligned anti-parallel or parallel MT cross-links plotted as the mean ± SD and compared with a two-way ANOVA. GX-WT, anti-parallel MTs n=54 and parallel MTs n=4; GX-MBD1^mut^, anti-parallel MTs n=47 and parallel MTs n=15; GX-MBD2^mut^, anti-parallel MTs n=40 and parallel MTs n=31. N=4 independent experiments. ns, not significant; *, p<0.05; **, p<0.01; ****, p<0.0001. (D-E) Weaker functional MBD1 (GX-MBD2^mut^) slides anti-parallel MTs faster than functional MBD2 (GX-MBD1^mut^) (D). Tighter functional MBD2 statically cross-links more parallel MTs than functional MBD1 (E). MT plus-ends are indicated by a plus sign (+) and the direction of MT sliding and motor motility are indicated by arrows.

If tail MT association inversely correlates with MT sliding (Braun *et al*., 2017), we might expect to see increased sliding velocity of MTs cross-linked with GX-MBD2^mut^, which has weaker tail binding to the MT. To investigate this idea, we measured the sliding velocity of aligned anti-parallel and parallel MT sliding events mediated by GX-WT, GX-MBD1^mut^, or GX-MBD2^mut^ (Figure 7A). For some cargo MTs, they either glided on the chamber surface before forming MT cross-links or slid off the MT cross-link and continued gliding (Figure 7A, GX-WT Anti-parallel Sliding, 4-10’). Because these gliding events had faster velocities (Figure S4A), we were careful to measure the velocities of only the sliding events in the aligned MT cross-links. Of the aligned sliding MTs that GX-WT slid, 93% were anti-parallel and had a mean velocity of 36 nm/s (Figure 7C), consistent with previous sliding velocities for GFP-XCTK2 at these concentrations (Hentrich and Surrey, 2010). GX-MBD1^mut^ slid 76% of the sliding MTs anti-parallel with a velocity similar to GX-WT, 33 nm/s. In contrast, GX-MBD2^mut^ slid 56% of the sliding MTs anti-parallel with a mean velocity of 60 nm/s that was approximately two-fold faster than GX-WT and GX-MBD1^mut^, p<0.0001 (Figure 7C). Despite GX-MBD2^mut^ sliding parallel MTs to a greater extent (Figure 7B), the sliding velocity of parallel MT cross-links was similar between the proteins with an average velocity of 33 nm/s, indistinguishable from the anti-parallel sliding velocities of GX-WT and GX-MBD1^mut^. These results provide evidence that tail MT affinity alone can modulate anti-parallel MT sliding velocity (Braun *et al*., 2017).

Some cargo MTs in our assay slid at angles across the lattice or across the ends of template MTs, which were mostly the minus ends (Figure S4B, top), or the cargo MT first slid at an angle to the template before becoming aligned (Figure S4B, bottom). These angled sliding events appeared faster than the sliding events of aligned MTs. It has been proposed that decreasing the ability of the tail to diffuse within a MT cross-link increases sliding velocity, which can manifest when cross-linked MTs slide at angles to one another (Lüdecke *et al*., 2018). To investigate whether the different affinities of the MT binding domains impacted this mechanism, we measured the sliding velocity of MTs that slid at an angle to the template and those that were aligning to the template MTs (Figure S4C and Video S4). GX-WT slid cargo MTs at an angle with a mean velocity of 58 nm/s, 61% faster than its aligned anti-parallel MT sliding of 36 nm/s, p<0.0001. GX-MBD1^mut^ slid angled cargo MTs with a similar velocity to GX-WT with a mean velocity of 61 nm/s, which was 91% faster than its aligned anti-parallel MT sliding of 32 nm/s, p<0.0001. In contrast, GX-MBD2^mut^ slid cargo at an angle with a mean velocity of 78 nm/s that was 30% faster than its aligned anti-parallel sliding of 60 nm/s, p<0.001, and 35% faster than GX-WT angled sliding velocity, p<0.0001, and 28% faster than GX-MBD1^mut^ angled MT sliding velocity of 61 nm/s, p<0.01. Consistent with the findings of Ncd angled MT sliding (Lüdecke *et al*., 2018), the sliding velocity of XCTK2 angled MT cross-links was faster than the sliding of aligned anti-parallel sliding MTs. However, the reduced MT affinity of GX-MBD2^mut^ increased the sliding velocity of angled MTs relative to GX-WT and GX-MBD1^mut^, suggesting that the affinity of the tail and diffusibility of the tail may modulate sliding velocity by different mechanisms. Together, our results suggest that the weaker MT affinity of MBD1 in the tail enables faster anti-parallel MT sliding velocity (Figure 7D) and the stronger MBD2 controls the extent of static parallel MT cross-linking (Figure 7E) for balanced robust anti-parallel MT sliding and tight parallel MT cross-links that contribute to efficient spindle assembly and morphogenesis.

## Discussion

The bipolar spindle is essential for accurate chromosome segregation, and molecular motors cross-link and slide MTs to sort and organize them into the classic fusiform shape of the spindle. K-14s are minus-end directed motors that cross-link and slide both anti-parallel and parallel MTs, but the mechanisms by which they cross-link and slide MTs in both orientations and how these activities contribute to spindle assembly are not clear. We show that the tail domain of *Xenopus* Kinesin-14, XCTK2, contains two MT binding domains with different MT affinities that regulate anti-parallel MT sliding velocity and the degree of parallel MT cross-linking, which contributes to proper spindle morphogenesis. These findings provide a mechanism for how K-14s balance different MT cross-linking and sliding activities and provide a framework for understanding how molecular motors regulate spindle assembly.

### K-14s contain two MT binding domains that differentially regulate anti-parallel MT sliding velocity and parallel MT cross-linking

It is well-established that kinesin tails are important for cargo binding and for the regulation of motor function (Yildiz, 2024). There are multiple mitotic kinesins that have an ATP-independent MT binding domain in their tail domain (Liao *et al*., 1994; Fink *et al*., 2009; Weaver *et al*., 2011; Weinger *et al*., 2011), but few have been shown to have more than one independent MT binding domain (Karabay and Walker, 1999; Shrestha *et al*., 2023). Previous work with *Drosophila* Kinesin-14 Ncd suggested that there are two regions in the tail that are involved in MT binding, which had different MT affinities (Karabay and Walker, 1999; Szczęsna and Kasprzak, 2016). In our study, we systematically identified two independent MT binding domains in the *Xenopus* XCTK2 tail, one within each half of the tail, and then used site-directed mutagenesis of K/R residues to disrupt MT binding of each MT binding domain individually. Analogous to the Ncd tail (Szczęsna and Kasprzak, 2016), the XCTK2 wild-type full-length tail had tight MT affinity in the sub micromolar range. However, the second MT binding domain in XCTK2, MBD2, had a four-fold higher MT affinity than MBD1, whereas the Ncd MT binding domains had opposite strengths in MT affinity in which the first MT binding domain bound MTs three-fold tighter than the second MT binding (Szczęsna and Kasprzak, 2016). The differences in MT affinities is not likely due to the organization of the domains within the tail because the first MT binding domain of Ncd also contains the NLS (Goshima and Vale, 2005), consistent with the XCTK2 tail (Ems-McClung *et al*., 2004). A potential caveat to comparing their MT affinities to ours is that in their studies the first MT binding domain was mutated by site-directed mutagenesis, whereas the second domain was deleted, which could impact overall protein structure (Szczęsna and Kasprzak, 2016). Our discovery that the XCTK2 tail also contains two MT binding domains of differing MT affinities suggests that this could be a conserved trait of the Kinesin-14 family. This concept is further supported by our unpublished work wherein the human HSET tail also contains two independent MT binding domains in each half of the tail, and like in Ncd, the N-tail appears to bind MTs more tightly (Cassity and Walczak, unpublished). The presence of two MT binding domains in the HSET tail is substantiated by the observation that the HSET tail displays bimodal rupture forces from MTs in optical trap assays that differ by three-fold (Reinemann *et al*., 2018), akin to the differences in MT affinities between the XCTK2 and Ncd tails. Together these studies favor the idea that Kinesin-14 tails, at least in the animal kingdom, contain two MT binding domains of differing MT affinities, and raises the question of how these domains are used to regulate MT cross-linking and sliding.

Multiple studies suggest that the ability of the Kinesin-14 tail to diffuse within a MT cross-link regulates its ability to slide anti-parallel MTs, but two different mechanisms are proposed to govern MT sliding. HSET and XCTK2 enrich within the overlap regions of anti-parallel cross-linked MTs, and as the density of the motors was increased, sliding velocity decreased (Hentrich and Surrey, 2010; Braun *et al*., 2017). These decreases were attributed to K-14 crowding within the overlap that slowed diffusion from the combined MT binding of both the motor and tail domains. Modeling suggested that either reduced binding of the motor domain or increased unbinding of the tail domain would increase sliding, which they demonstrated by increasing the ionic strength of the buffer to reduce tail MT binding (Braun *et al*., 2017). Our previous work with XCTK2 in a higher ionic strength buffer showed that anti-parallel MTs slid 41% faster than parallel MTs (Ems-McClung *et al*., 2020) compared to our current study in lower ionic strength buffer where they had similar sliding velocities. These results are consistent with the idea that reduced MT affinity of K-14s allows for faster MT sliding. Alternatively, it was suggested that reducing the ability of the Ncd tail to diffuse within a MT cross-link increases sliding velocity because Ncd slid anti-parallel MTs three-fold slower than MTs that slid at angles to one another, presumably as a result of restricting tail diffusion to a small region (Lüdecke *et al*., 2018). We also found that GX-WT angled MT sliding velocity was about two-fold faster than aligned anti-parallel sliding MT velocity. However, our finding that the weaker GX-MBD2^mut^ also slid these angled MTs faster than GX-WT and GX-MBD1^mut^ indicates that the affinity of the tail is still a mechanism for regulating sliding velocity in angled MT sliding. One interpretation of the findings by Lüdecke (Lüdecke *et al*., 2018) could be that in the angled crossovers, there are fewer motors allowing for more diffusion rather than restricting diffusion, which would be consistent with the inverse correlation of sliding velocity and motor density observed by others (Hentrich and Surrey, 2010; Braun *et al*., 2017). Our findings that there are two Kinesin-14 tail MT binding domains with different MT affinities that can independently regulate sliding may help reconcile these results.

While several studies have addressed how K-14s slide anti-parallel MTs, much less is known about how K-14s statically cross-link parallel MTs. An attractive model for the function of the two K-14 MT binding domains was that each MT binding domain acted in one type of MT cross-link, anti-parallel or parallel. However, our finding that both tail MT binding domains cross-link both MT orientations to similar proportions goes against this idea. We did find that the weaker MT affinity of MBD1 resulted in more parallel MT sliding, and the tighter MBD2 had more static parallel cross-linking, but there was no difference in parallel MT sliding velocities. Consistent with our current results with the weaker GX-MBD2^mut^, our previous MT sliding assays with XCTK2 in higher ionic strength buffer resulted in similar numbers of sliding and statically cross-linked parallel MTs (Ems-McClung *et al*., 2020). These results suggest that the affinity of the tail may play a role in the extent of parallel MT sliding but not the velocity of sliding, implying that modulation of anti-parallel and parallel sliding velocities may have different mechanisms. Additional work will need to be done to reconcile the differences between these various studies. For example, how does changing the affinity of a single MT binding domain affect the proportion and velocity of MTs that slide within MT cross-links, and how are those changes correlated with the ability of the tail to diffuse and with the density of the motors within the cross-links? In addition, can both MT binding domains engage with the MT at one time, or is there only a single MT binding domain engaged during active MT sliding?

### Kinesin-14 tail mediated MT cross-linking and sliding differentially contributes to spindle localization and morphogenesis

K-14s are important for spindle assembly, for spindle length control, and for focusing MTs at poles, all activities that likely require MT cross-linking and sliding. The amount of K-14s on the spindle also appears to be an important factor as over addition of XCTK2 to egg extracts stimulates spindle assembly (Walczak *et al*., 1997) as does mild overexpression of HSET in oocytes (Bennabi *et al*., 2018); however, high HSET overexpression in oocytes causes spindle collapse (Bennabi *et al*., 2018). In addition, increasing the levels of RanGTP in the spindle results in an increase in the localization of endogenous XCTK2 to the spindle (Ems-McClung *et al*., 2020), suggesting that localization is tightly regulated. How the amount of K-14s on the spindle and the differential MT cross-linking and sliding of XCTK2 contribute to spindle assembly and morphogenesis is unclear. Our findings suggest that tight parallel MT cross-linking is important for efficient spindle assembly because GX-WT and GX-MBD1^mut^ over addition stimulated spindle formation, whereas GX-MBD2^mut^ did not stimulate spindle assembly to the same extent. However, a caveat to these findings is that GX-MBD2^mut^ did not localize well to spindles. One possibility for the reduced localization of GX-MBD2^mut^ is that it has reduced MT binding in MgADP+P_i_ compared to GX-WT and GX-MBD1^mut^, indicating that the tighter MBD2 MT binding may be important for the maintenance of XCTK2 on the spindle during its ATPase cycle. It is also possible that different mechanisms or interacting partners of the tail domains control the turnover of XCTK2 within the spindle, resulting in differential localization. For example, we observed previously that inhibiting importin binding by mutations in the NLS of XCTK2 increased its turnover in the spindle, and that the turnover of XCTK2 was differentially regulated in the spindle midzone and at the poles (Weaver *et al*., 2015), implying that interaction with the importins spatially regulates XCTK2 MT association. Consistent with this idea, our line scan analysis suggests that both MT binding domains are needed for robust association of GX-WT on the spindle as GX-MBD1^mut^ and GX-MBD2^mut^ had overall reduced levels of spindle association, although the reduction in GX-MBD1^mut^ localization did not significantly hinder its ability to stimulate spindle assembly. These results suggest that it will be important to understand not only how the differential MT affinities of the Kinesin-14 tail domains impact their dynamic localization on the spindle, but also how they contribute to efficient spindle assembly.

MT organization in spindles from different organisms as well as between meiosis and mitosis can vary, which suggests that the types of MT cross-linking and sliding activities may not be fully conserved. For example, the long spindle phenotype from overexpression of K-14s in mammalian mitotic cells (Matuliene *et al*., 1999; Cai *et al*., 2009) might mean that MBD1 in mammalian cells has slower anti-parallel sliding due to higher MT affinity, or that there is a switch in the strength of the MT binding domain affinities, as seen with the Ncd tail (Szczęsna and Kasprzak, 2016). In support of this idea, over addition of wild-type HSET to *Xenopus* egg extracts increased spindle length relative to control whereas over addition of a similar amount of WT XCTK2 did not (Cai *et al*., 2009), suggesting an intrinsic feature of HSET was important for the increase in spindle length. Alternatively, the observed differences in effects of K-14s on spindle length in mitotic cells could be a consequence of the differences in MT organization in *Xenopus* spindles, which are meiotic in origin. Meiotic spindle assembly in *Xenopus* is driven by chromatin-mediated MT nucleation, which results in a dense array of MTs in the center of the spindle attached to two polar arrays on either side (Yang *et al*., 2008; Houghtaling *et al*., 2009) versus human mitotic spindles that have more MTs emanated from the centrosome with less anti-parallel overlap in the midzone (Nixon *et al*., 2017; O’Toole *et al*., 2020). Consistent with this idea, our line scan analysis of GX-WT addition indicates an increase in MT polymer in the middle of the spindle, suggesting either increased nucleation or stabilization of MTs in this region. It is also possible that both the tail MT binding domain affinities and cell type contribute to these various phenotypes. Our work highlights the need for future experiments to better understand how the Kinesin-14 MT binding domains assist in spindle assembly in different systems, with a focus on how they contribute to the dynamics of spindle MT organization.

In addition to the ability of the Kinesin-14 MT binding domains to impact the efficiency of spindle assembly, they also contributed to spindle morphology. The addition of either GX-WT or GX-MBD1^mut^ to *Xenopus* spindle assembly assays resulted in spindles with narrow widths, which correlates with their better ability to statically cross-link parallel MTs. Overexpression of mammalian K-14s in cells also results in narrow spindles (Matuliene *et al*., 1999; Cai *et al*., 2009). GX-MBD2^mut^, which did not statically cross-link parallel MTs as well as GX-WT and GX-MBD1^mut^, had shorter spindles, which may mean that the faster anti-parallel sliding of GX-MBD2^mut^ could antagonize the outward forces of Kinesin-5 and/or Kinesin-12 mediated anti-parallel sliding. This antagonistic relationship has been well-described in the literature for spindles in multiple systems (Saunders *et al*., 1997; Mountain *et al*., 1999; Sharp *et al*., 2000; Yukawa *et al*., 2018) as well as *in vitro* reconstitutions (Peterman and Scholey, 2009; Hentrich and Surrey, 2010; Reinemann *et al*., 2018). It is interesting that the shortened spindle morphology that occurs in the presence of excess GX-MBD2^mut^ is contrary to the long spindle morphology we observed when a vast excess of a partial MBD1 mutant (NLSb) was added to *Xenopus* spindle assembly reactions (Cai *et al*., 2009). In that study, we proposed that the addition of excess NLSb mutant lengthened spindles because of increased parallel sliding, but our current findings suggest a more complicated scenario. Because the NLSb mutation reduces the MT affinity and capacity of MBD1, this may mean that in the full-length protein, NLSb also inhibited or reduced MBD1 sliding activities like GX-MBD1^mut^ in addition to retaining high static parallel MT cross-linking. The absence or reduction of the faster inward MBD1-mediated anti-parallel sliding in the midzone due to the NLSb mutation could allow for more outward parallel sliding that could lengthen the spindle. However, this long spindle phenotype is only seen with a vast excess of the NLSb mutant, suggesting it may be a gain of function phenotype. Perhaps the increased static parallel MT cross-linking with high levels of NLSb results in the elongation of the tiled MTs in *Xenopus* spindles (Yang *et al*., 2007), which would be consistent with the extremely thin and wispy appearance of the MTs in those spindles (Cai *et al*., 2009). Together, our data suggests that the parallel cross-linking activity of MBD2 balances the more robust MBD1 mediated anti-parallel sliding needed for spindle length control.

### Mechanisms that regulate Kinesin-14 mediated MT cross-linking and sliding

K-14s interact with various proteins through their N-terminal tail domain, like CEP215 (Chavali *et al*., 2016) and NuMA (Cutillas and Johnston, 2021), which may govern pole focusing. Kinesin-14 tails also interact with the plus-tip binding protein, EB1 (Braun *et al*., 2013), which is proposed to regulate MT plus end dynamics by removing EB1 from the plus tips, resulting in increased catastrophes (Ogren *et al*., 2022). Alternatively, it has been suggested that EB1 may provide a hand hold for K-14s to bind to growing MT plus ends where it helps align newly nucleated MTs into parallel arrays (Molodtsov *et al*., 2016), which are needed for kinetochore clustering and biorientation in yeast (Kornakov *et al*., 2020). Interestingly, robust EB1-mediated Ncd plus-tip tracking required at least one tail MT binding domain, suggesting the need for Kinesin-14 tails to make a MT contact when complexed with EB1 (Szczęsna and Kasprzak, 2016). How the MT binding domains contribute to EB1-mediated tip tracking in other K-14s and how these interactions are differentially regulated is unknown.

One master regulator in the spindle is the RanGTP gradient, which has high RanGTP around the chromatin that diminishes toward the spindle poles (Kaláb *et al*., 1999; Wilde and Zheng, 1999; Kaláb *et al*., 2002; Kaláb *et al*., 2006). The Ran effectors, importin α/β, bind to spindle assembly factors containing an NLS, which can result in a spatial effector gradient response in the spindle (Ems-McClung *et al*., 2020). The gradient itself is important in modulating Kinesin-14 MT association such that turnover at the chromatin is slower than at the poles (Weaver *et al*., 2015). Our current finding that the tail contains two MT binding domains with one containing an NLS supports the idea that these domains may be differentially regulated by the RanGTP gradient. One possibility is that the importins inhibit MBD1 from binding MTs near the poles, an idea consistent with our previous findings using fluorescence lifetime imaging microscopy (FLIM) showing that the importins interact with XCTK2 through the NLS within the spindle near the poles (Ems-McClung *et al*., 2020) and with our data showing that the turnover of GFP-XCTK2 is faster near the poles than near the spindle equator (Weaver *et al*., 2015). In this model MBD2, whose MT binding is stronger and is not directly regulated by the importins, could readily bind to the parallel MTs to focus and stabilize them. Thus, K-14s having two independent MT binding domains with different binding affinities may allow the RanGTP gradient to spatially control Kinesin-14 function. In addition to regulation by Ran, phosphorylation of Kinesin-14 tails may provide additional control. Kar3-Cik1 is phosphorylated by Cdk1, which inhibits EB1 interaction (Molodtsov *et al*., 2016), and phosphorylation at S26 in the HSET EB1 binding site by ATM and ATR kinases promotes centrosome clustering in cancer cells (Fan *et al*., 2021). Ncd is proposed to be regulated by Aurora B phosphorylation at S96, which is in the NLS of the MT binding domain, and this phosphorylation promotes tail-MT interaction near the chromatin (Beaven *et al*., 2017). Mutation of the NLS in XCTK2 reduced the ability of Aurora B to phosphorylate XCTK2 *in vitro* (Zhang and Walczak, unpublished), suggesting Aurora B phosphorylation may regulate XCTK2 activity as well. Together these studies raise the question about the regulatory mechanisms that direct the spatial and temporal control of K-14 association with its binding partners, which in turn governs spindle morphogenesis.

### How is the tail domain structured to mediate interactions with multiple binding partners

Understanding how the small tail domain of K-14s (12-20 kDa depending on the organism) mediates interactions with multiple binding partners is perplexing. Structural predictions using AlphaFold (Abramson *et al*., 2024) indicate that K-14 tails are highly disordered. This may be advantageous to allow these regions to adopt multiple conformations depending on the binding partner and/or to have the flexibility to bind MTs simultaneously with other binding partners in temporally and spatially regulated modes. Our previous FLIM results show that the importins bind XCTK2 near the poles (Ems-McClung *et al*., 2020), which could be mediated through the motor domain only or could be that MBD2 binds to MTs when MBD1 is bound to importins. Our current MT binding assays show that the importins fully inhibit the binding of the YPet-Tail N proteins, which only contain MBD1, but that the importins only partially inhibit binding of the full YPet-Tail, which contains both MBD1 and MBD2. These results support the idea that MBD2 might bind MTs while MBD1 is bound to importins. Consistent with this idea, structural modeling with ColabFold predicts that when MBD1 is bound to importin α/β the rest of the tail is sufficiently disordered that MBD2 would be available to bind MTs. However, in our MT pelleting assays, we do not find stoichiometric co-sedimentation of the importins with YPet-Tail, which would suggest that the YPet-Tail bound to MTs does not have importins bound. This result supports another idea that while the importins inhibit MBD1 from binding MTs, they cannot simultaneously be bound to MBD1 when MBD2 is bound to MTs. If this model is correct, then our FLIM results would suggest that the importin bound XCTK2 is bound to MTs through the motor domain. Given that GX-MBD1^mut^, which cannot interact with MTs through MBD1, enriches at the poles in our spindle assembly assays like GX-WT, indicates that a population of XCTK2 bound to MTs through MBD2 exists in addition to XCTK2 bound to importins.

Modeling also predicts that each MT binding domain interacts with either α- or β-tubulin, which is consistent with our affinity assays that show each MT binding domain has saturable binding with tubulin monomer. Further support for this idea comes from the studies showing that MT binding of the HSET tail (Reinemann *et al*., 2018) and the Ncd tail (Karabay and Walker, 2003) were differentially sensitive to the removal of either the α-tubulin or both the α- and β-tubulin C-terminal tails, implying that there may be one MT binding site on the lattice and one site involving the C-terminal tails of tubulin. However, these binding interactions may be more complex as an early cryoEM study found that the Ncd tail made contacts with four sites on tubulin dimer, two on each monomer (Wendt *et al*., 2003). In the future it will be critical to identify these MT interaction sites at the higher resolution offered by the explosion in EM technology (Nogales and Scheres, 2015; Nogales and Mahamid, 2024), which will provide additional insights into how the differential interaction with the MT governs the ability of K-14s to cross-link and slide both parallel and anti-parallel MTs in the spindle. When combined with studies of how these different MT binding domains regulate the spatial organization of MTs in the spindle, we will gain new insights into how regulated interactions in the spindle direct proper spindle morphogenesis and function.

## Materials and Methods

### Cloning and mutagenesis

To generate the N-terminal deletion series of the XCTK2 tail domain fused to YPet, cDNAs were amplified by PCR from pRSETA-YPet-XCTK2-Tail (amino acids 2-120) (Ems-McClung *et al*., 2020) or pRSETA-YPet-24GNM that contains the tail and stalk domains (non-motor domain, NM) (amino acids 2-289) as shown in Table 1 using the primers listed in Table 2. pRSETA-YPet-24GNM, was generated by amplifying YPet from pRSETA-Rango-2 (Kaláb *et al*., 2002; Kaláb and Soderholm, 2010) using the GFP*BamH*IF1 and GFP*SacI*R3 primers (Table 2) and replacing mCitrine in pRSETA-mCitrine-24GNM using *BamH*I/*Sac*I restriction enzymes (Table 1). Plasmid pRSETA-mCitrine-24GNM was generated by subcloning the cDNA encoding the non-motor domain of XCTK2 (amino acids 2-289) from pFastBac1-GFPXCTK2 (Cai *et al*., 2009) into pRSETA-mCitrine (Ems-McClung *et al*., 2013) using the 5’*Sac*I24G and XCTK2-NM-*Kpn*I-R primers (Table 2). For the C-terminal tail domain deletion series tagged with either YPet or CyPet, cDNA was amplified from pRSETB-24GNM, pRSETA-YPet-24GNM, or pRSETA-YPet-XCTK2-Tail and cloned according to Table 1 using the primers shown in Table 2. The cloning vector for the CyPet tagged Tail C2, CyPet-Tail C2, plasmid pRSETA-CyPet, was generated by amplifying the DNA encoding CyPet from pRSETA-Rango-2 with the GFP*BamH*IF1 and GFP*Sac*IR5 primers (Table 2), digesting with *BamH*I/*Sac*I, and cloning into pRSETA. All clones were verified by sequencing.

**Table 1.** Expression plasmids used in this study.

| *Expression Plasmid* | *Template or Insert* | *Forward Primer* | *Reverse Primer* | *Parent Plasmid* |
| --- | --- | --- | --- | --- |
| pRSETA-YPet-24GNM | pRSETA-Rango-2 | GFPBamHIF1 | GFPSacIR3 | pRSETA-mCitrine-24GNM |
| pRSETA-YPet-XCTK2-Tail | pRSETA-YPet-24GNM | T7 Forward | XCTK2-A120-stop-HindIIIR | pRSETB-5'YPet |
| pRSETA-YPet-Tail N1 | pRSETA-YPet-24GNM | T7 Forward | XCTK2-S53-stop-HindIIIR | pRSETB-5'YPet |
| pRSETA-YPet-Tail N2 | pRSETA-YPet-XCTK2-Tail | 5'SacI24G | XCTK2-A61-stop-HindIIIR | pRSETA-YPet-24GNM |
| pRSETA-YPet-Tail N3 | pRSETA-YPet-XCTK2-Tail | 5'SacI24G | XCTK2-A79-stop-HindIIIR | pRSETA-YPet-24GNM |
| pRSETB-YPet-Tail C1 | pRSETA-YPet-24GNM | XCTK2-I54-SacIF | XCTK2-A120-stop-HindIIIR | pRSETB-5'YPet |
| pRSETB-YPet-Tail C2 | pRSETA-YPet-XCTK2-Tail | XCTK2-A62-SacIF | XCTK2-A120-stop-HindIIIR | pRSETB-5'YPet |
| pRSETB-YPet-Tail C3 | pRSETA-YPet-XCTK2-Tail | XCTK2-V80-SacIF | XCTK2-A120-stop-HindIIIR | pRSETB-5'YPet |
| pRSETA-CyPet-Tail C2 | pRSETA-YPet-XCTK2-Tail | XCTK2-A62-SacIF | XCTK2-A120-stop-HindIIIR | pRSETA-5'CyPet |
| pRSETA-YPet-Tail N3-NLSb | pRSETA-YPet-XCTK2-Tail-NLSb | 5'SacI24G | XCTK2-A79-stop-HindIIIR | pRSETA-YPet-24GNM |
| pRSETA-YPet-Tail N3-MBD1<sup>mut</sup> | pRSETB-24GNM-NLSab/MBD1<sup>mut</sup> | 5'SacI24G | XCTK2-A79-stop-HindIIIR | pRSETA-YPet-24GNM |
| pRSETA-5'CyPet | pRSETA-Rango-2 | GFPBamHIF1 | GFPSacIR5 | pRSETA |
| pRSETA-CyPet-Tail C2-NLS3 | pRSETB-24GNM-NLS3 | XCTK2-A62-SacIF | XCTK2-A120-stop-HindIIIR | pRSETA-5'CyPet |
| pRSETA-CyPet-Tail C2-NLS3-Mut3 | pRSETA-CyPet-Tail C2-NLS3 | XCTK2Mut3F | XCTK2Mut3R | SDM |
| pRSETA-CyPet-Tail C2-MBD2<sup>mut</sup> (NLS3-Mut3-Mut2) | pRSETA-CyPet-Tail C2-NLS3-Mut3 | XCTK2Mut2F | XCTK2Mut2R | SDM |
| pRSETA-YPet-Tail-MBD1<sup>mut</sup> | pRSETB-24GNM-NLSab/MBD1<sup>mut</sup> | 5'SacI24G | XCTK2-A120-stop-HindIIIR | pRSETB-5'YPet |
| pRSETA-YPet-Tail-NLS3 | pRSETB-24GNM-NLS3 | 5'SacI24G | XCTK2-A120-NLS3-stop-HindIIIR | pRSETB-5'YPet |
| pRSETA-YPet-Tail-MBD2<sup>mut</sup> | pRSETA-YPet-Tail-NLS3 | XCTK2Mut2F, XCTK2Mut3F | XCTK2Mut2R, XCTK2Mut3R | Sequential SDM |
| pRSETA-YPet-Tail-MBD1<sup>mut</sup>-NLS3 | pRSETB-CyPet-Tail C2-NLS3 | RD: Sall/HindIII |  | pRSETA-YPet-Tail-MBD1<sup>mut</sup> |
| pRSETA-YPet-Tail-MBD1-2<sup>mut</sup> | pRSETA-YPet-Tail-MBD1<sup>mut</sup>-NLS3 | XCTK2Mut2F, XCTK2Mut3F | XCTK2Mut2R, XCTK2Mut3R | Sequential SDM |
| pFastBac1-GFP-XCTK2 | pFB-GFP-XCTK2 |  |  |  |
| pFBGFP-XCTK2-MBD1<sup>mut</sup> | pGEX1*-24GNM-NLSab | 5'SacI24G | pGEX3' | pFBGFP-XCTK2 |
| pFBGFP-XCTK2-MBD2<sup>mut</sup> | pRSETB-24GNM-MBD2<sup>mut</sup> | RD: Sall/Spel |  | pFBGFP-XCTK2 |
| pFBGFP-XCTK2-MBD1-2<sup>mut</sup> | pRSETB-24GNM-MBD2<sup>mut</sup> | RD: Sall/Spel |  | pFBGFP-XCTK2-MBD1<sup>mut</sup> |
| pFastBac1-6His-GFP | p6HisGFP | 5'6HGFP | GFPSacIR2 | pFastBac1-GFP |
| pFB6HGFP-XCTK2-MBD2<sup>mut</sup> | pFBGFP-XCTK2-MBD2<sup>mut</sup> | RD: SacI/KpnI |  | pFB6HGFP |
| pFB6HGFP-XCTK2-MBD1-2<sup>mut</sup> | pRSETB-24GNM-MBD2<sup>mut</sup> | RD: Sall/Spel |  | pFBGFP-XCTK2-MBD1<sup>mut</sup> |

SDM: Site-directed mutagenesis. RD: Restriction digest.

**Table 2.**
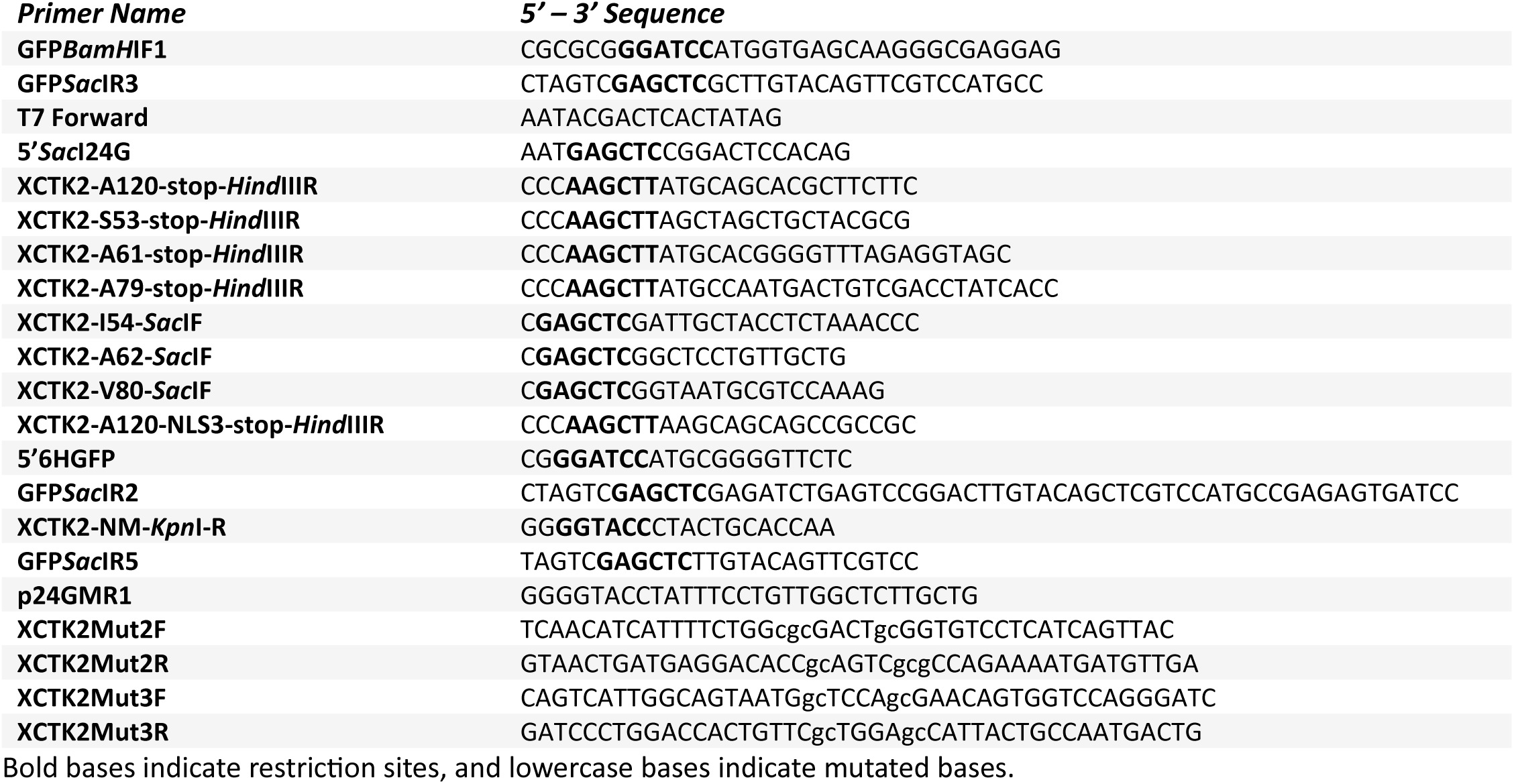
Cloning and mutagenesis primers used to generate expression plasmids.

The cDNAs of the XCTK2 tail containing the NLS mutations that disrupt MBD1 (NLSab/MBD1^mut^) were amplified by PCR from pRSETB-24GNM-NLSab/MBD1^mut^ (Ems-McClung *et al*., 2004) and cloned into pRSETA-YPet-24GNM according to the conditions in Tables 1 and 2 to generate YPet-Tail N3-NLSb and YPet-Tail N3-MBD1^mut^ constructs. The mutations that disrupt MBD2 (NLS3, Mut2, and/or Mut3) were generated by first creating pRSETA-CyPet-Tail C2-NLS3 from pRSETB-24GNM-NLS3 (Ems-McClung *et al*., 2004), and then sequentially mutating pRSETA-CyPet-Tail C2-NLS3 (Table 1) using the QuikChange^TM^ system with the XCTK2Mut2F/R and XCTK2Mut3F/R mutagenesis primers (Table 2). To create the full-length pRSETA-YPet-Tail with the MBD1 ^mut^, the corresponding cDNA was amplified from pRSETB-24GNM-NLSab/MBD1^mut^ and cloned as described above for pRSETA-YPet-Tail using the conditions in Tables 1 and 2. The pRSETA-YPet-Tail-MBD2^mut^ expression plasmid was created by sequential site-directed mutagenesis on pRSETA-YPet-Tail-NLS3 using conditions in Tables 1 and 2. The MBD double mutant tail expression plasmid, pRSETA-YPet-Tail-MBD1-2^mut^, was generated by subcloning the NLS3 mutation in pRSETB-CyPet-Tail C2-NLS3 into pRSETA-YPet-Tail-MBD1^mut^ with subsequent sequential site-directed mutagenesis as described in Tables 1 and 2. All clones were verified by sequencing.

For baculovirus production, full-length XCTK2 constructs containing the MBD1^mut^, MBD2^mut^, or MBD1-2^mut^ mutations were generated by subcloning the corresponding cDNA regions from either pGEX1*-24GNM-NLSab (Ems-McClung *et al*., 2004) or pRSETB-24GNM-MBD2^mut^ into pFastBac1-GFP-XCTK2 (Cai *et al*., 2009) *Sac*I/*Spe*I restriction sites to create pFBGFP-XCTK2-MBD1^mut^ or pFBGFP-XCTK2-MBD2^mut^ or by subcloning from pRSETB-24GNM-MBD2^mut^ into pFBGFP-XCTK2-MBD1^mut^ *Sal*I/*Spe*I to create pFBGFP-XCTK2-MBD1-2^mut^ using the conditions in Table 1 and 2. Plasmids for pFB6HGFP-XCTK2-MBD2^mut^ and pFB6HGFP-XCTK2-MBD1-2^mut^ were made by subcloning the cDNA of XCTK2 from pFBGFP-XCTK2-MBD2^mut^ or pFBGFP-XCTK2-MBD1-2^mut^ into the *Sac*I/*Kpn*I sites of pFastBac1-6His-GFP. The pFastBac1-6His-GFP vector was constructed by subcloning the 6His-GFP DNA sequence from p6HisGFP (Walczak *et al*., 2002) into the *BamH*I/*Sac*I sites of pFastBac1-GFP (Hertzer *et al*., 2006). All clones were verified by sequencing.

### Expression and purification of recombinant proteins

For bacterial expression of the YPet- or CyPet-tagged Tail, N- and C-terminal tail deletion proteins, and corresponding mutants (Table 1), BL21(DE3)pLysS bacteria harboring the corresponding expression plasmids, were induced and purified using Ni-NTA agarose (Qiagen, #30210) as described previously for the purification of YPet-XCTK2-Tail (Ems-McClung *et al*., 2020). In brief, cells were grown in 1 L LB containing 50 μg/ml ampicillin, 34 μg/ml chloramphenicol, with or without 2 mM MgSO_4_ at 37°C and induced for the expression of recombinant proteins with 0.1 mM isopropyl β-D-1-thiogalactopyranoside at 22°C for 24 h. Cells were pelleted, snap frozen in liquid nitrogen, and stored at −80°C. For protein purification, the cells were briefly thawed and lysed on ice in Lysis Buffer (50 mM phosphate buffer, pH 8.0, 300 mM NaCl, 10 mM imidazole, 0.1% Tween-20, 1 mM phenylmethylsulfonyl fluoride (PMSF), 1 mM benzamidine, 1 μg/ml LPC (1 μg/ml leupeptin, 1 μg/ml pepstatin, 1 μg/ml chymostatin), 0.5 mg/ml lysozyme) by sonication using a Branson Sonifier 250 four to six times at 20% output. Following sonication, the NaCl concentration was increased to 500 mM and β-mercaptoethanol (β-ME) was added to a final concentration of 5 mM. The crude lysate was spun at 18,000 rpm for 20 min at 4°C in a Beckman JA25.5 rotor. The supernatant (cleared lysate) was added to Ni-NTA agarose equilibrated with Column Buffer (50 mM phosphate pH 8.0, 500 mM NaCl, 0.1% Tween-20, 10 mM imidazole) and incubated for 1 h at 4°C with rotation. The lysate-agarose mixture was centrifuged in a clinical centrifuge for 4 min at 1,500 rpm to pellet the beads with bound protein. The beads were washed twice in 10 column volumes (CV) of Column Buffer containing 1 mM PMSF, 1 mM benzamidine, 1 μg/ml LPC, and 5 mM β-ME for 10 min with rotation at 4°C. Ten CV of Column Wash Buffer (50 mM phosphate buffer, pH 6.0, 500 mM NaCl, 5 mM β-ME, 0.1 mM PMSF, 1 μg/ml LPC) was added to the beads and loaded into a Bio-Rad Poly-Prep® Chromatography Column (#7311550) and allowed to drain by gravity flow at 4°C. The bound protein was eluted with 10 CV of Elution Buffer (50 mM phosphate buffer, pH 7.2, 500 mM NaCl, 400 mM imidazole, 5 mM β-ME, 1 μg/ml LPC) at 4°C in 1 ml aliquots. Aliquots with the highest fluorescence were dialyzed into 1 L of XB250 dialysis buffer (10 mM HEPES, pH 7.2, 250 mM KCl, 50 mM sucrose, 0.1 mM EDTA, and 0.1 mM EGTA) at 4°C overnight followed by dialysis with an addition 1 L of XB250. The dialyzed protein was clarified at 14,000 rpm at 4°C for 15 minutes, snap-frozen in liquid nitrogen in single use aliquots and stored at −80°C. The expression and purification of His_6_-GFP, His_6_-importin α-CyPet, importin α-His_6_, and S-His_6_-importin β were performed similarly with dialysis into XB dialysis buffer (10 mM HEPES, pH 7.2, 100 mM KCl, 25 mM NaCl, 50 mM sucrose, 0.1 mM EDTA, and 0.1 mM EGTA) (Ems-McClung *et al*., 2004; Ems-McClung *et al*., 2020).

For baculoviral expression of full-length GFP-tagged wild-type XCTK2 and the MBD mutants, baculovirus stocks were produced according to the Bac-to-Bac system (Gibco^TM^, #10359016) as previously described (Cai *et al*., 2009). Briefly, pFastBac1-GFP-XCTK2 wild-type and mutant plasmids were transposed into DH10Bac bacmid cells and screened by blue/white colony selection and PCR. Sf9 cells were cultured in SF-900 II SFM media (Invitrogen) + 0.5X Antibiotic-antimycotic (Life Technologies, #15240062) at 27°C in shaking flasks. Sf9 cells were transfected with ∼1 μg bacmid DNA using CellFectin™ II Reagent (Invitrogen, #10362-100) according to the manufacturer’s protocol. After 72-96 h when >90% of cells were transfected based on GFP fluorescence, the media containing virus was collected (Pass 0), centrifuged at 500 x g for 5 min, removed to a fresh tube, supplemented with 5% FBS, and stored at 4°C wrapped in aluminum foil. Pass I and Pass II viral stocks were made by infecting ∼2 x 10^6^ cell/ml with Pass 0 or Pass I virus at a multiplicity of infection (MOI) of 0.1 for 72 h at 27°C with shaking. Media containing virus particles was harvested by pelleting the cells at 500 x g for 5 min, filtering the media, supplementing with 5% FBS, and storing at 4°C wrapped in aluminum foil. For protein expression in Sf9 cells, 400-600 ml of ∼2 x 10^6^ cell/ml were infected with Pass II virus at a MOI of 2-5 for 42-72 h. Cells were harvested by centrifuging at 1,000 x g for 5 min at 4°C, the cell pellet was frozen in liquid nitrogen and stored at −80°C.

GFP-XCTK2 and GFP-XCTK2-MBD1^mut^ proteins were purified as described previously (Walczak *et al*., 1997). Briefly, cells were lysed in 20-40 ml of BRB80 (80 mM PIPES pH 6.8, 1 mM MgCl_2_, 1 mM EGTA), 1 mM dithiothreitol (DTT), 250 mM KCl, 0.1 mM EDTA, 2 mM MgATP, 1 mM PMSF, 1 μg/ml LPC on ice with sonication. The clarified lysate was filtered through an AcroPak 20 0.8/0.2 μm Filter with Supor Membrane (Pall Corporation, #12202) and applied onto a 1 ml HiTrap SP column (Cytiva, #17-1151-01) equilibrated in 100 mM KCl, FPLC buffer (20 mM PIPES pH 6.8, 1 mM MgCl_2_, 1 mM EGTA, 0.1 mM EDTA, 1 mM DTT, 10 μM MgATP, 0.1 μg/ml LPC) attached to an ÄKTA Pure and controlled by Unicorn software (Cytiva). Protein was eluted with a linear 100 mM – 1 M KCl gradient in 1 ml fractions. Fractions with the highest GFP intensity were pooled, filtered with a 0.2 μm filter, applied in 1 ml fractions to a 24 ml Superose 6 Increase 10/300 GL (Cytiva, #29-0915-96) column equilibrated in 30% FPLC buffer (FPLC buffer containing 300 mM KCl), and eluted in 0.5 ml fractions. The fractions with the highest GFP intensity were pooled, concentrated in an Amicon® Ultra-0.5 Centrifugal Filter Concentrator with Ultracel® 100 Regenerated Cellulose Membrane NMWL:100,000 (EMD Millipore, #UFC510024) if needed, solid sucrose added to 10%, flash frozen in single use aliquots in liquid nitrogen, and stored at −80°C.

For purification of 6HisGFP-XCTK2-MBD2^mut^ and 6HisGFP-XCTK2-MBD1-2^mut^ proteins, Sf-9 cell pellets were lysed in 80 mM PIPES pH 6.8, 2 mM MgCl_2_, 300 mM KCl, 10 mM imidazole, 2 mM ATP, 5 mM β-ME, 1 mM PMSF, 1 μg/ml LPC and sonicated with a Branson Sonifier 250 two times for 20 seconds at 20% output. The lysate was clarified by centrifugation at 18,000 rpm in a JA25.5 rotor for 20 min. The clarified soluble lysate was incubated with Ni-NTA agarose equilibrated in the lysis buffer for 1 h at 4°C with rotation. The agarose beads were washed once in 10 CV lysis buffer and once in 10 CV wash buffer (20 mM PIPES pH 6.8, 1 mM MgCl_2_, 300 mM KCl, 10 mM imidazole, 10 µM ATP, 5 mM β-ME, 0.1 μg/ml LPC). Bound protein was eluted in 1 ml fractions with 10 ml elution buffer (20 mM PIPES pH6.8, 1 mM MgCl_2_, 300 mM KCl, 400 mM imidazole, 10 µM ATP, 5 mM β-ME, 0.1 μg/ml LPC). Fractions with the highest GFP intensity were pooled, and ∼2 mL of sample was buffer exchanged using a 5 ml 7K molecular weight cut off Zeba™ Spin Desalting Column (ThermoFisher, #89891) equilibrated in 30% FPLC buffer according to the manufacturer’s directions and then filtered with a 0.2 μm filter. The filtered Ni-NTA purified protein was applied in 1 ml fractions to a 24 ml Superose 6 10/30 column equilibrated in 30% FPLC buffer and eluted in 0.5 ml fractions. The fractions with the highest GFP intensity were pooled, concentrated in an Amicon® Ultra-0.5 Filter, solid sucrose added to 10%, flash frozen in single use aliquots in liquid nitrogen, and stored at −80°C. All proteins were quantified with the Bradford Assay followed by densitometry of the full-length proteins of sodium dodecyl sulfate polyacrylamide gel electrophoresis (SDS-PAGE) gels stained with colloidal Coomassie Blue using BSA as a control. To assess the purity of the recombinant proteins, 0.5 ug of each protein was electrophoresed on a 10% SDS-PAGE gel and stained with InstantBlue® Coomassie Protein Stain (ISB1L) (Abcam, #ab119211).

### MT binding assays

MTs were polymerized from 10 μM cycled bovine tubulin in BRB80/DTT (BRB80, 1 mM DTT) with 0.5 mM guanylyl- (α,β)-methylene-diphosphonate (GMPCPP; Jena Bioscience, #NU-405L) and 20 μM paclitaxel at 37°C for 30 min to generate doubly stabilized MTs. MTs were sedimented at 45,000 rpm in a TLA100 rotor for 12 min at 35°C in a Beckman Optima MAX-TL centrifuge and resuspended in BRB10/DTT/paclitaxel (BRB10 (10 mM PIPES pH 6.8, 1 mM MgCl_2_, 1 mM EGTA), 1 mM DTT, 10 μM paclitaxel). MT concentration was determined by the A_280_ of tubulin dimer using 115,000 M^-1^cm^-1^ as the molar extinction coefficient.

Bulk MT binding assays were performed by mixing 5 μM MTs in BRB10/DTT/paclitaxel with 0.5 μM YPet-tagged XCTK2 tail proteins with or without 2 μM importin α-His_6_ and S-His_6_-importin β in 10 mM HEPES pH 7.7, 25 mM KCl, 50 mM sucrose, 0.1 mM EDTA, 0.1 mM EGTA, 0.2 μg/μl casein. Proteins were incubated for 15 min and sedimented at 45,000 rpm for 12 min at 22°C in a TLA100 rotor. The supernatant was removed, and the pellet resuspended in an equal volume of buffer. Equivalent volumes of the supernatants and pellets were electrophoresed on 10% SDS-PAGE gels and stained with colloidal Coomassie Blue. The fraction bound was determined by densitometry of the full-length proteins and then corrected for background pelleting of a no MT control and multiplied by 0.5 μM to convert to amount protein bound. The average protein bound was determined and plotted as the mean ± SD from at least three independent experiments. YPet-Tail, YPet-Tail-MBD1^mut^, and YPet-Tail-MBD2^mut^ with and without importins were compared using a two-way ANOVA with Tukey’s multiple comparisons test in GraphPad Prism. YPet-Tail and YPet-Tail-MBD1-2^mut^ without importins were compared using a Student’s *t*-test in Prism. Significance was considered if the *p* value was less than 0.05.

Bulk MT binding assays in MgATP or MgADP+P_i_ were performed with 0.1 μM full-length GFP-XCTK2 or MBD mutants in the absence or presence of 0.4 μM importin α-His_6_ and S-His_6_-importin β and 1 μM MTs similar to those done with the YPet-Tail proteins except that the reaction buffer consisted of 20 mM PIPES pH 6.8, 75 mM KCl, 1 mM MgCl_2_, 1 mM EGTA, 0.1 mM EDTA, 0.5 μg/μl casein and either 2 mM MgATP or 2 mM MgADP + 20 mM P_i_. The supernatant and resuspended pellet fractions were added to a Nunc 384-well plate (#262260), and the GFP fluorescence measured by exciting at 470 nm and reading the emission at 510 nm in a BioTek Synergy H1 monochromator-based plate reader (Ems-McClung and Walczak, 2020). The average protein bound was determined in Excel and plotted as the mean ± SD from at least three independent experiments and compared with a three-way ANOVA with Tukey’s multiple comparisons test in Prism. Significance was considered if the p value was less than 0.05.

### Apparent affinity assays

Importin α/β affinity assays using Förster resonance energy transfer (FRET) (Ems-McClung and Walczak, 2020) were performed with 0.1 μM His_6_-importin α-CyPet/0.4 μM S-His_6_-importin β and 0-2 μM YPet-Tail, YPet-Tail N3, YPet-Tail C2, or YPet-Tail C3 in 10 mM HEPES pH 7.7, 25 mM KCl, 50 mM sucrose, 0.1 mM EDTA, 0.1 mM EGTA, 1 mM DTT, 0.2 μg/μl casein. Samples were excited at 405 nm and detected with a spectral emission scan from 440-600 nm using 5 nm steps in a Synergy H1 plate reader. Spectral emissions were corrected for YPet emission bleed through by subtracting a control reaction without importin α-CyPet/importin β at each YPet-Tail protein concentration. The YPet emission at 530 nm, the peak emission of YPet, was normalized to the CyPet emission at 460 nm to determine the FRET ratio. The FRET ratios were normalized to 100% fraction bound for a FRET ratio equal to 5, the maximum FRET observed with YPet-Tail and then adjusted to micromolar importin α-CyPet bound by multiplying by 0.1 μM. The amount of importin α-CyPet/importin β bound was fit with a quadratic equation for binding (Foster *et al*., 1998) and graphed as the mean ± SD from 4-7 independent experiments. Differences in apparent importin affinity compared to YPet-Tail were determined using the extra sum-of-squares F-test for variance in Prism in which the *K*_d_ and *B*_max_ were shared.

MT affinity assays were performed analogous to the bulk MT assay except that the reaction buffer contained 0.5 μg/μl casein, and the MT concentration in terms of tubulin dimer ranged from 0-10 μM. The fraction bound was determined by densitometry of the full-length tail proteins from colloidal Coomassie Blue stained gels and normalized in Prism between 0 and 1 to account for non-specific pelleting of the proteins. The normalized fraction bound was converted to the amount of tail protein bound by multiplying by 0.5 μM, the concentration of tail protein in the reaction. The mean protein bound ± SD from at least three independent experiments was determined and graphed in Prism with the best fit curve. The wild-type YPet-Tail binding to MTs fit the stoichiometric binding curve (Bell *et al*., 2017) with a stoichiometry coefficient of 2.4, indicating the tail bound to two sites on tubulin dimer, which could correspond to one site on each tubulin monomer. The YPet-Tail MT binding was then fit to the quadratic equation for saturable binding (Foster *et al*., 1998) in terms of tubulin monomer, which fit well based on the replicates test for lack of fit and the graph of the residuals. For comparison of apparent MT affinities, all the tail proteins were plotted as a function of tubulin monomer in the main figures. Significance between apparent affinity curves was determined using the extra sum-of-squares F-test for variance in Prism in which the *K*_d_ and *B*_max_ were both shared.

### Spindle assembly and immunofluorescence

Cytostatic factor (CSF) extracts were made from *Xenopus laevis* eggs (Murray, 1991) using cytochalasin B (cyto B, Sigma-Aldrich, #C6762) in place of cytochalasin D. CSF extract was supplemented with 1/1000 volume of 10 mg/ml LPC, 1/1000 volume of cyto B, 1/50 volume of 50x energy mix (375 mM creatine phosphate, 50 mM ATP, 5 mM EGTA, 50 mM MgCl_2_), 1/40 volume of 2M sucrose, and 0.4 μM Biotium X-rhodamine (VWR, #90005) labeled tubulin or 0.14 µM Pierce™ NHS-rhodamine (Thermo Scientific, #46406) labeled tubulin. For CSF spindle assembly, sperm was added to 250 sperm/µl and 50 nM (monomer concentration) GFP, GFP-XCTK2 wild-type, or mutant proteins were added and incubated at RT for 60 min. Cycled spindles were assembled from CSF extracts supplemented with 500 sperm/µl and 10x calcium chloride solution (10 mM HEPES pH7.7, 1 mM MgCl_2_, 100 mM KCl, 150 mM sucrose, 10 µg/ml cyto B, and 4 mM CaCl_2_), and incubated at room temperature until the sperm nuclei entered interphase (∼80 min) followed by addition of an equal volume of CSF extract. 50 nM GFP, GFP-XCTK2, or mutant proteins (monomer) were added, and the extracts were incubated for 60 min for bipolar spindle assembly. For immunofluorescence, reactions were fixed in 2% formaldehyde, 2.5 mM MgCl_2_, 30% glycerol, 0.5% Triton X-100, BRB80, layered on top of a 40% glycerol/BRB80 cushion, and sedimented onto coverslips. Coverslips were post-fixed in cold methanol for 5 min, rehydrated in TBS-Tx (10 mM Tris pH 7.6, 150 mM NaCl, 0.1% Triton X-100), and DNA stained with 10 μg/ml Höechst 33258 (Sigma-Aldrich, #14530) in TBS-Tx. Coverslips were mounted in Prolong Diamond™ Antifade Mountant (Life Technologies, #P36961), cured for ∼24 h, and sealed with nail polish.

Each coverslip was imaged as a 3×3 (CSF) or 10×10 (cycled) montage on a Nikon NiE microscope equipped with a Nikon 40X Plan Apo 1.0 NA objective, Hamamatsu ORCA-FLASH Monochrome 30FPS camera, and Lumencore Spectra III Light Engine light source controlled by Nikon NIS-Elements software. For spindles assembled by CSF assembly, GFP fluorescence was detected with the 475/28 nm bandpass filter at 1% power for 100 ms, MTs were detected with the 555/28 nm filter at 20% power for 500 ms (X-rhodamine tubulin), and DNA with the 390/22 nm filter at 15% power for 100 ms (Höechst). For the percent spindle assembly in CSF extracts, ∼100 structures (asters, half spindles, spindles) were counted per experiment from five to six independent experiments, plotted in Prism with the mean ± SD, and analyzed by an ordinary one-way ANOVA with Tukey’s multiple comparisons test. For spindles assembled in cycled extracts, GFP fluorescence was detected at 20% or 31.9% power for 100 ms, MTs at 35% power for 300 ms (rhodamine tubulin), and DNA at 15% power for 100 ms (Höechst). Images were background-subtracted using a rolling ball algorithm set to 35.08 μm for the GFP and MT channels, and 8.03 μm for the DNA channel using NIS-Elements General Analysis 3 software. For the representative images, each channel was scaled the same using Fiji, and images were assembled in Adobe Illustrator. Cycled spindles that were isolated from other structures in the image and not overlapping the edge of the image were outlined using the NIS-Elements “Simple ROI Editor” and “Auto Detect” features and converted to binary objects. For each binary spindle object, the integrated MT intensity, integrated GFP intensity, max Feret diameter (length), min Feret diameter (width), and area were exported to Excel, plotted in Prism with the mean ± SD, and analyzed by a Kruskal-Wallis test with Dunn’s multiple comparisons test.

To measure the distribution of MTs and the GFP-XCTK2 proteins across the spindles, line scans were performed on rolling ball background subtracted cycled spindles by manually drawing a 15 pixel width line from pole-to-pole and normalized to 101 bins across the length of the spindle using the XLineScan plugin for Fiji (Wilbur and Heald, 2013).

The normalized MT integrated intensity, GFP integrated intensity, and ratio of GFP to MT integrated intensity were plotted as the mean ± SEM in Prism. To calculate the difference in protein distribution across the spindle between the proteins, the ratios of GFP to MT intensity were compared using a Mann-Whitney test, and the *p* values were plotted in Prism. Note that the significant *p* values in the body of the spindle between GX-WT and GX-MBD1^mut^ were between *p* < 0.05 and *p* < 0.001, and for GX-WT vs GX-MBD2^mut^ and GX-MBD1^mut^ vs GX-MBD2^mut^ the values all along the spindle were below *p* < 0.0001.

For Western analysis, extract was diluted 1:10 into 2X SB (125 mM Tris pH 6.8, 4% SDS, 20% glycerol (w/v), 4% β-ME, bromophenol blue), heated at 100°C for 5 min, and electrophoresed on a 10% SDS-PAGE gel. Protein was transferred to BioTrace™ NT nitrocellulose (Pall Corporation, #66485), probed with 0.5 μg/ml rabbit anti-XCTK2-NM (Walczak *et al*., 1997) for XCTK2 and 1:10,000 mouse DM1α (Sigma-Aldrich, #T9026) for tubulin followed by 1:5,000 donkey anti-rabbit or 1:10,000 sheep anti-mouse HRP linked whole secondary antibodies, developed with SuperSignal™ West Pico PLUS Chemiluminescent Substrate (Thermo Scientific, #34577), and visualized using a Bio-Rad ChemiDoc™ Imaging System. Digital images were assembled using Adobe Photoshop and Illustrator. For each Western (CSF or cycled), the GFP-XCTK2 (GX) and tubulin (Tub) loading control bands came from the same blot.

### MT cross-linking assays

Polarity marked MTs were assembled from X-rhodamine or HiLyte647 (Cytoskeleton, #TL670M-A) labeled tubulin seed and elongation mixes, each containing 0.5 mM GMPCPP in BRB80, 1 mM DTT. The mixes were clarified by centrifugation at 45,000 rpm for 7 min at 2 °C in a TLA100 rotor, frozen in liquid nitrogen in single use aliquots, and stored at −80 °C. Short MT seeds were polymerized at 1-2 μM tubulin in BRB80/DTT for 60 min. MT extensions were then polymerized from the freshly made seeds by diluting 1:10 into 0.5 μM or 2 μM elongation mix and incubated at 37 °C for 60 min. The polarity marked MTs were sedimented at 45,000 rpm for 12 min at 35 °C in a TLA100 rotor and resuspended in BRB80/1 mM DTT/10 μM paclitaxel. MT concentration was determined by A_280_ as described above for MTs. Template MTs were 8% X-rhodamine-tubulin and 8% biotin-tubulin (Cytoskeleton, #T333P) labeled seeds with 4% HiLyte647- and 46% biotin-labeled extensions. Cargo MTs were composed of 9% HiLyte647 labeled seeds with 13% X-rhodamine labeled extensions.

To visualize MT sliding, flow chambers (∼10 μl volume) were assembled on a slide with double-stick tape using biotin-polyethylene glycol (Laysan Bio, #SVA-5000) coated 22 x 30 mm and 22 x 22 #1.5 coverslips (Ems-McClung *et al*., 2013; Ems-McClung *et al*., 2020). Chambers were rinsed consecutively with two volumes of BRB80/DTT and 5% Pluronic® F-127 (Sigma, #P2443) in BRB80, and incubated for 3 min. The chamber was rinsed with two volumes of Block (20 mM PIPES pH6.8, 75 mM KCl, 1 mM MgCl_2_, 0.1 mM EDTA, 1 mM EGTA, 1 mM DTT, 1 mg/ml casein, 10 μM paclitaxel), and then rinsed and incubated with two volumes of 0.05 mg/ml NeutrAvidin® (Life Technologies, #A-2666) diluted in Block for 3 min. After rinsing with Block, 0.4 μM template MTs in Block were introduced into the chamber and incubated for 3 min. The chamber was rinsed with two volumes of Block and then 5 nM GFP-XCTK2 (GX-WT), 10 nM GX-MBD1^mut^, 10 nM GX-MBD2^mut^, or 10 nM GX-MBD1-2^mut^ in Block was introduced to the chamber, and incubated for 3 min. After rinsing with Block, 0.2 μM cargo MTs were introduced to the chamber and incubated for 5 min. All incubations were done at room temperature with the slide upside down. Sliding was induced by rinsing the unbound cargo MTs from the chamber with two volumes of Block + 2 mM MgATP supplemented with oxygen scavenging mix (0.32 mg/ml glucose oxidase, 0.055 mg/ml catalase, 5 mM DTT, 25 mM glucose). Chambers were imaged on a Nikon NiE equipped with a Nikon 60x Plan Apo 1.4 NA objective, Hamamatsu ORCA-FLASH Monochrome 30FPS camera, and Lumencore Spectra III Light Engine light source controlled by Nikon NIS-Elements software. Timelapse imaging was performed at 30 sec intervals for 10 min using a 3×3 grid. GFP-XCTK2 fluorescence was detected with the 475/28 nm bandpass filter at 1% power for 100 ms, and labeled segmented MTs were detected with the 555/28 nm filter at 15% power for 200 ms (X-rhodamine) and the 635/22 nm filter at 0.5% power for 100 ms (HiLyte647). The resulting nine movies from the 3×3 grid image were split and analyzed independently.

The first three movies (FOV) from four independent experiments for each of the protein conditions were scored for the number of cargo MTs present during the 10 min imaging time, and the numbers plotted as the mean ± SD and compared by a one-way ANOVA with Tukey’s multiple comparisons test in Prism. Because GX-WT cross-linked more MTs to the template MTs than GX-MBD1^mut^ or GX-MBD2^mut^ per movie, different numbers of movies were scored per condition to give 35-38 cross-linked MTs on average per condition per experiment. From these cross-linked MTs, the orientations of the MT cross-links were determined by switching between the X-rhodamine and HiLyte647 channels and comparing the brightnesses and lengths of the seeds and extensions under the assumption that plus ends had longer extensions than minus ends due to the differences in dynamic instability for plus and minus ends of MTs (Kristofferson *et al*., 1986). The percentage of aligned anti-parallel and parallel MTs was determined, plotted as the mean ± SD, and compared by a two-way ANOVA with Tukey’s multiple comparisons test in Prism. Cargo MTs were considered to slide if they slide for two or more frames (1 min), and cargo MTs were scored as static if they did not slide at all during the 10 min movie or did not slide after forming a MT cross-link. A single cargo MT could undergo multiple sliding events on different template MTs by sliding off one template onto another or by gliding in between template MTs, although this was infrequent. The percentage of aligned anti-parallel or parallel MTs that slid or were statically cross-linked was determined, plotted as the mean ± SD, and compared by a two-way ANOVA with Tukey’s multiple comparisons test in Prism. Velocities of aligned or angled sliding cargo events were measured using the Manual Tracking plugin in Fiji, exported to Excel where the average velocity was determined for each event, plotted in Prism as the mean ± SD, and compared with a two-way ANOVA with Tukey’s multiple comparisons test. Anti-parallel or parallel aligned sliding velocities were measured during the time frames when cargo MTs were aligned completely along the template axis. Gliding events were defined by cargo moving on the slide for two or more frames (1 min) independent of interactions with template MTs. Gliding event velocities were plotted with the aligned anti-parallel sliding velocities in Prism as the mean ± SD and compared by a two-way ANOVA with Tukey’s multiple comparisons test. Angled sliding cargo MTs either 1) slid across the ends of template MTs at an angle to the long axis of the template MT, which was most often the minus end, 2) slid at an angle across the lattice of the template MT, or 3) consisted of the time frame in which the cargo MT was at an angle to the template MT before becoming fully aligned. Angled sliding velocities were plotted as the mean ± SD and compared to the aligned anti-parallel velocities by a two-way ANOVA with Tukey’s multiple comparisons test in Prism. Representative images and movies were scaled the same in each channel. Movies were smoothed and annotated using Fiji and then converted from AVI to MP4 using Microsoft Clipchamp.

## Acknowledgments

This work was funded by NIH R35GM122482 (CEW) and by an NSF Transitions to Excellence Award (MCB 2128166) to CEW. MC was supported by a Cox Research Scholar Award. We thank Stephanie Zheng for help with the deletion analysis and bulk MT binding assays, Sanjay Shrestha for help in the mutagenesis, cloning, and expression of the YPet- and CyPet-Tail deletion series, and Lesley Weaver for the generation of cloning plasmids for full-length GFP-XCTK2. We also thank Sanjay Shrestha and Molly Lusk for careful and critical editing of the manuscript.

## Author Contributions

M. Cassity and S. Ems-McClung performed the YPet-Tail deletion analysis and bulk MT binding (Figure 1, B and C). S. Ems-McClung performed the importin α/β and MT affinity assays (Figures 1D-E, S2D, 2B-C, and 3B) and the YPet-Tail and GFP-XCTK2 bulk MT binding assays (Figure 3, C and D). M. Cassity cloned the NLS mutations into YPet-Tail N3 and CyPet-Tail C2 (Figure 2A). S. Ems-McClung performed and analyzed the CSF spindle assembly assays and corresponding Western (Figures 4 and S3A). A. Prasannajith performed the cycled spindle assembly assays, corresponding Western, and line scan analysis (Figures 5A, D-I, and S3B). S. Ems-McClung analyzed the cycled spindle assembly assays (Figures 5B-C, J-K, and S3C). S. Ems-McClung performed and analyzed the MT cross-linking assays (Figures 6, 7, and S4). S. Ems-McClung assembled all the figures and videos. C. Walczak oversaw the research, S. Ems-McClung and C. Walczak wrote the manuscript, and A. Prasannajith and M. Cassity helped edit the manuscript.

## Abbreviations

GX: GFP-XCTK2
HSET: human SET protein homolog
K-14: Kinesin-14
MBD: microtubule binding domain
MT: microtubule
Ncd: *Drosophila* non-claret disjunctional protein
NLS: nuclear localization signal
XCTK2: *Xenopus* carboxy-terminal kinesin 2

## Supplemental Figure Legends

**Figure S1.**
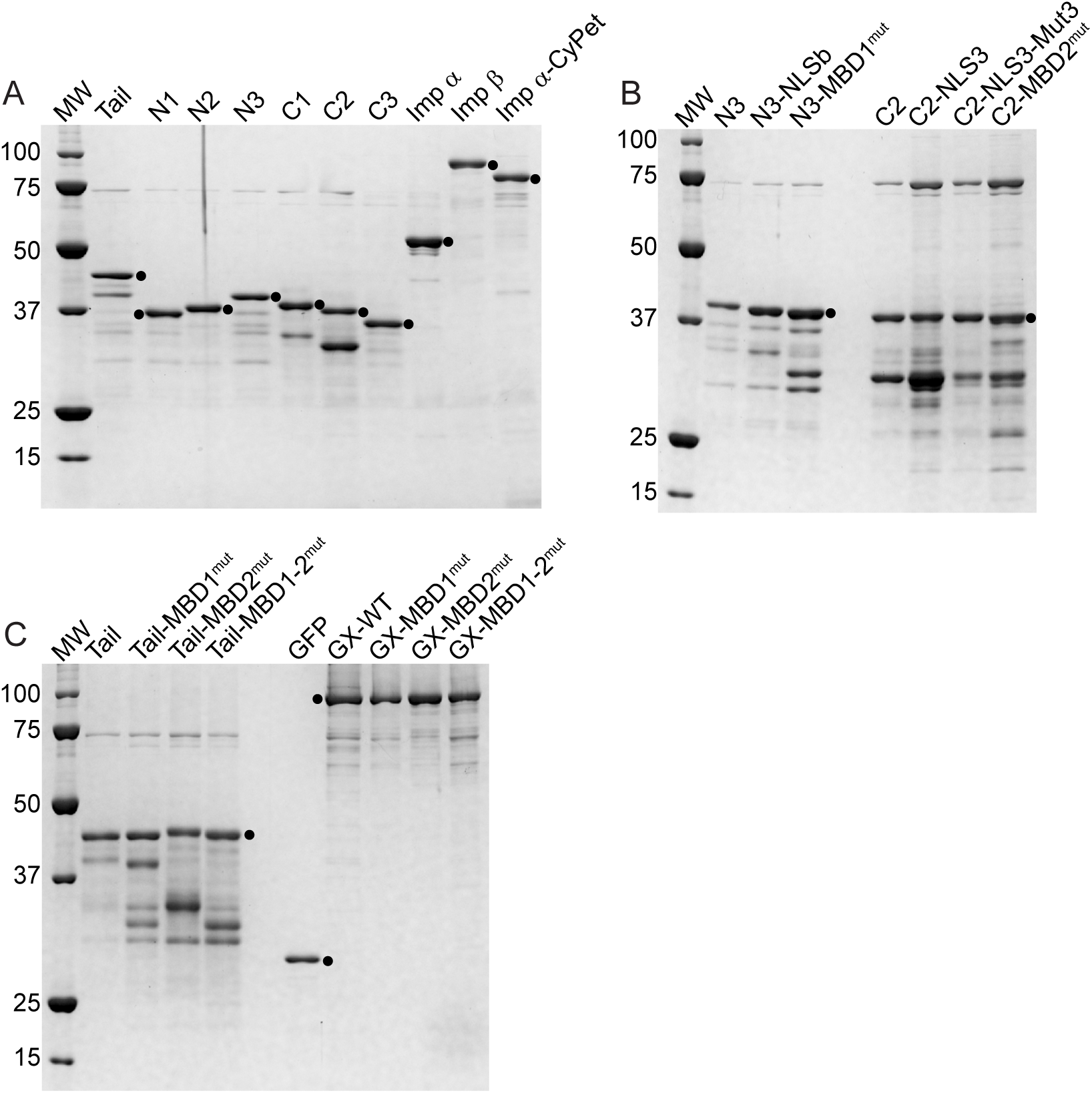
(A-C) Purified recombinant proteins (0.5 μg) used in the study were electrophoresed on 10% SDS-PAGE gels and stained with InstantBlue® Coomassie Protein Stain. (A) Proteins used in Figure 1, (B) proteins used in Figure 2, and (C) proteins used in Figures 3-7. The corresponding full-length proteins are indicated by a dot (●), and the molecular weight markers (MW) are Bio-Rad Precision Plus Protein ™ Unstained Protein Standards: 100, 75, 50, 37, 25, 20 and 15 kDa. Note that for all the MT binding and affinity measurements, the indicated molar concentrations are that of only the full-length protein and not of any truncated fragments or contaminants.

**Figure S2.**
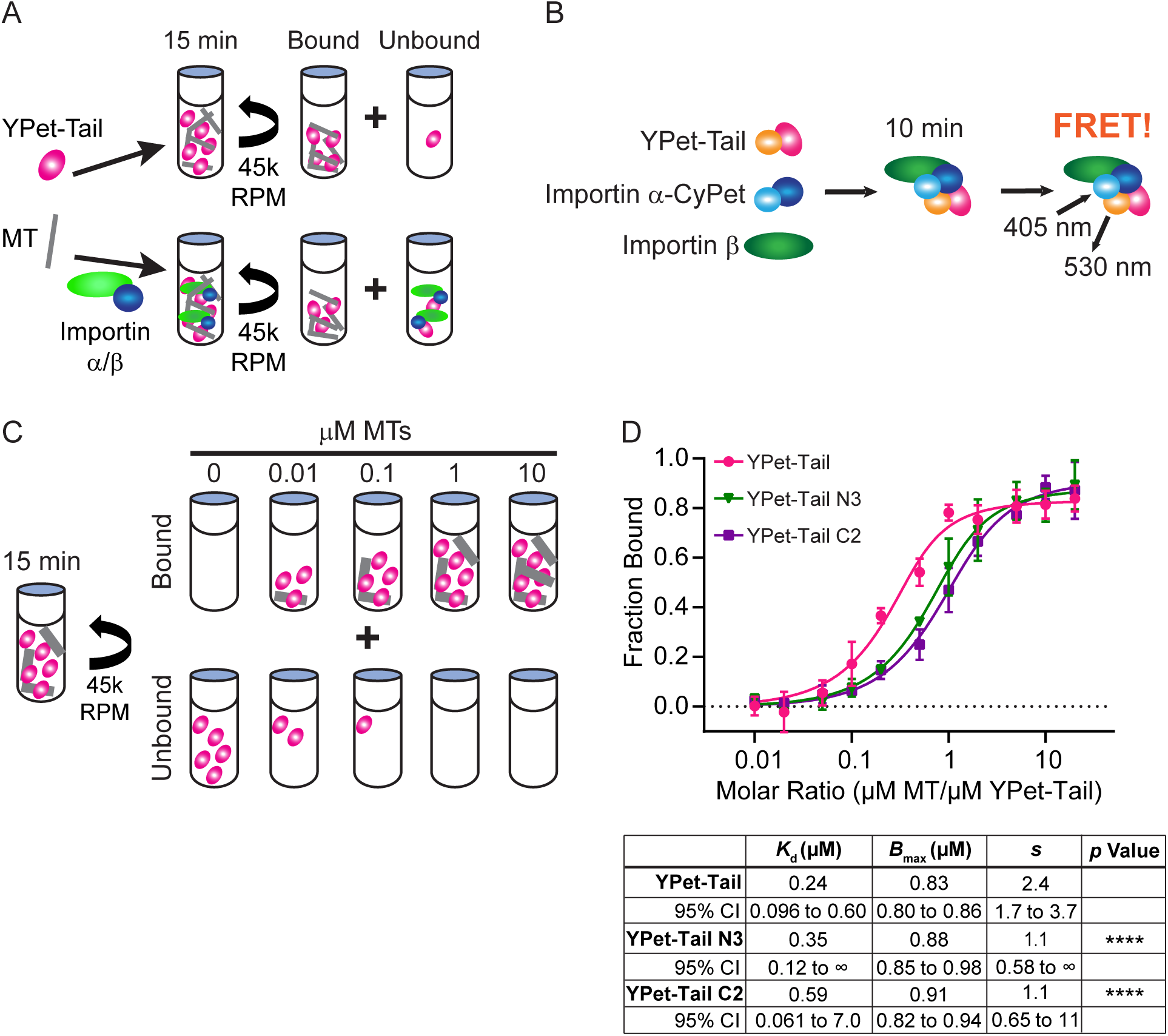
(A) Schematic of MT binding assays ± importin α/β using MT co-sedimentation. (B) Schematic of importin α-CyPet/importin β affinity assays with YPet-Tail using Förster resonance energy transfer. (C) Schematic of MT co-sedimentation assay to measure MT affinity. (D) Apparent MT affinity curves of 0.5 μM YPet-Tail, N3, or C2 graphed as the mean fraction ± SD of tail protein bound to increasing molar ratios of MTs to YPet-Tail protein and fit to the stoichiometry quadratic equation for saturable binding where the stoichiometry coefficient *s* represents the number of binding sites on a tubulin dimer. The apparent stoichiometric affinities between YPet-Tail and N3 or C2 proteins were compared with an extra sum-of-squares F-test. N=3-8 independent experiments. ****, p<0.0001.

**Figure S3.**
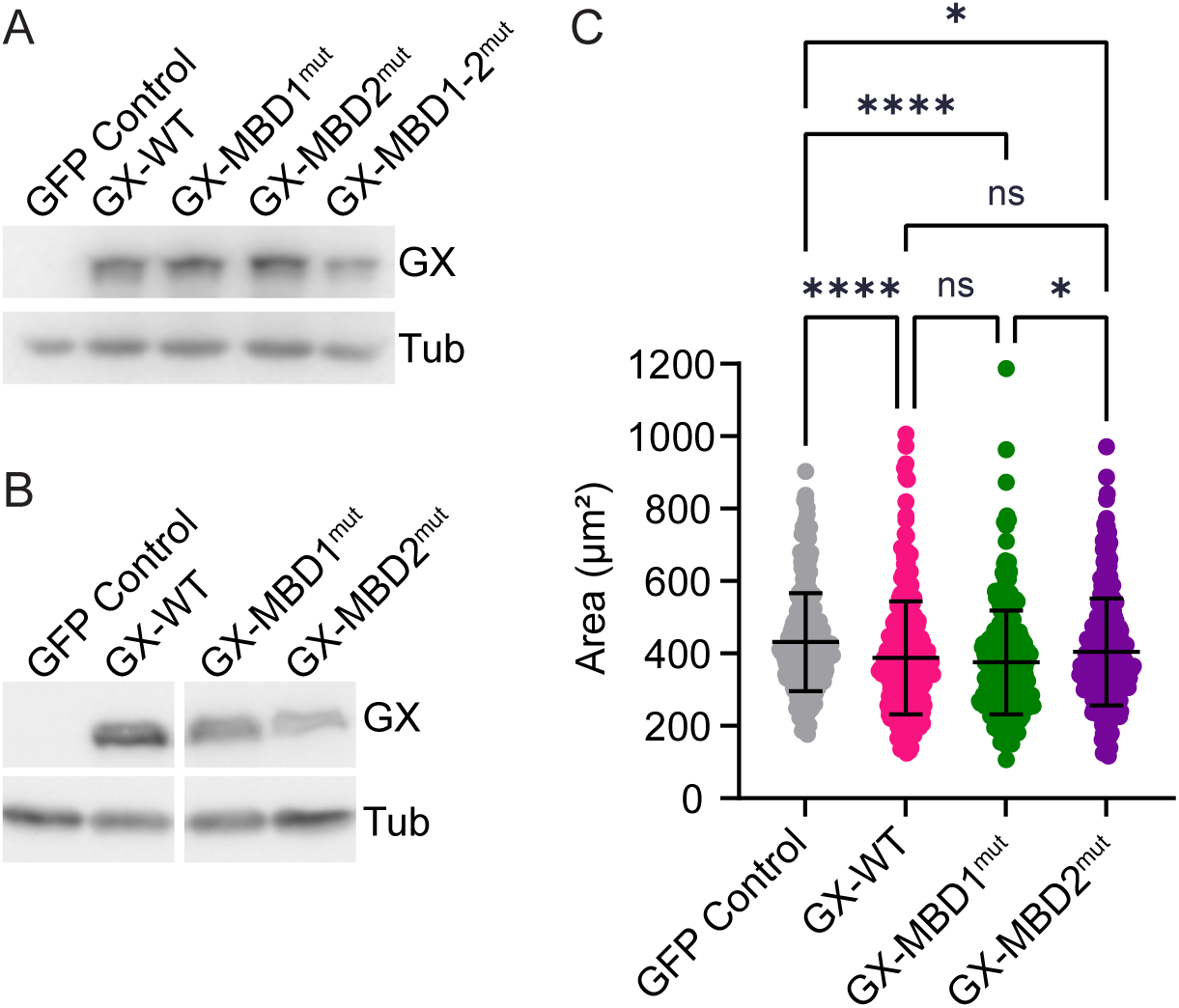
Western blots of GFP control, GX-WT, GX-MBD1^mut^, GX-MBD2^mut^, and/or GX-MBD1-2^mut^ addition to CSF (A) or cycled (B) spindle assembly reactions and probed with antibodies against XCTK2 (GX) and tubulin (Tub). Per Western, each panel is from the same blot. (C) Measured areas of cycled spindles plotted with the mean ± SD from N=3 independent experiments. Spindle numbers: GFP control n=274, GX-WT n=296, GX-MBD1^mut^ n=308, GX-MBD2^mut^ n=301. ns, not significant; *, p<0.05; ****, p<0.0001.

**Figure S4.**
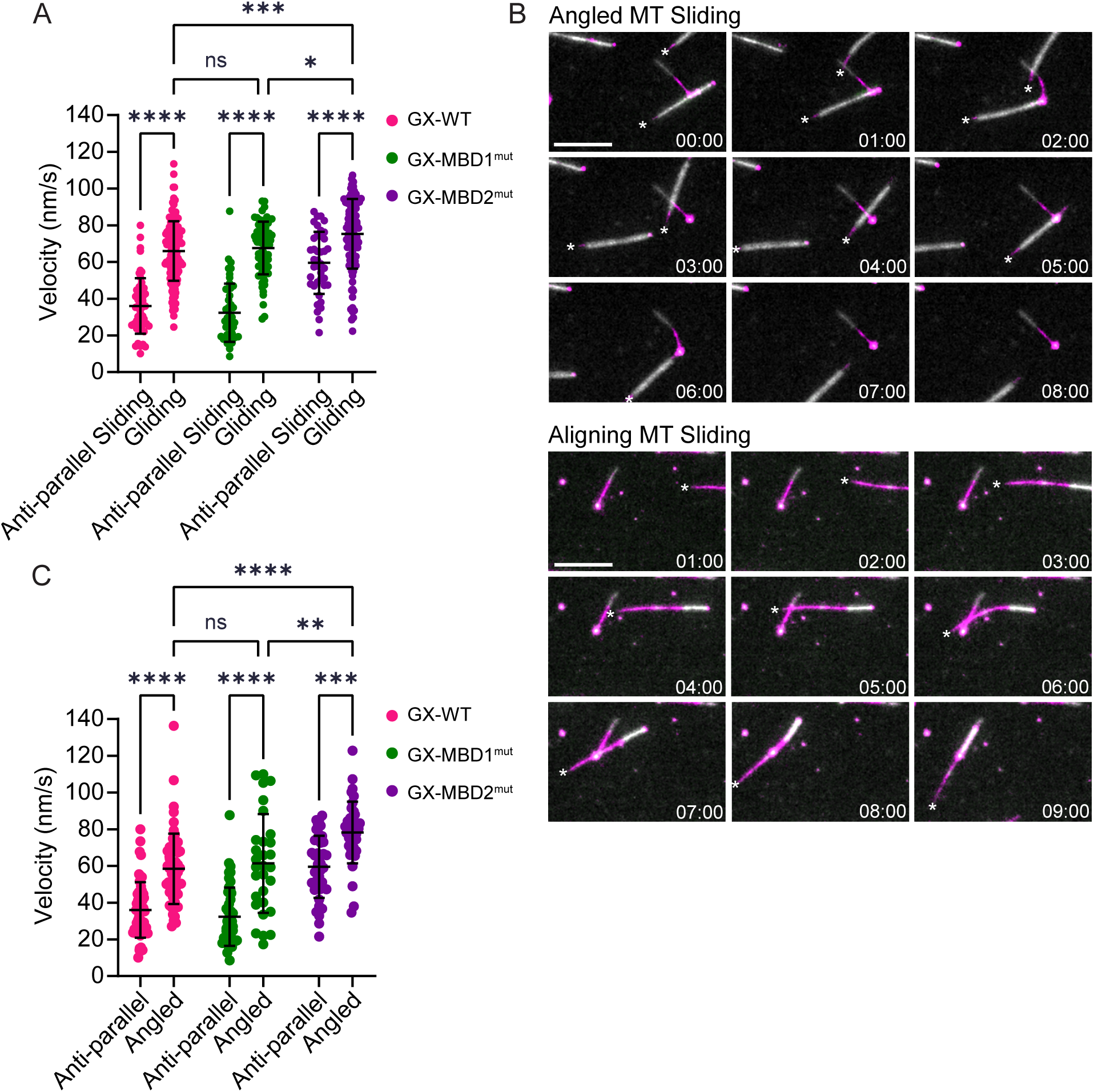
(A) Aligned anti-parallel sliding and gliding velocities plotted as the mean ± SD and compared with a two-way ANOVA. Anti-parallel sliding velocities are the same as plotted in Figure 7C. Gliding events are the motility of the MTs on the coverslip independent of the velocity of sliding in the MT cross-links: GX-WT, n=154; GX-MBD1^mut^, n=70; GX-MBD2^mut^, n=111 where n= the total number of MTs scored from N=4 independent experiments per condition. (B) Representative image series of angled (top) or aligning (bottom) MT sliding events from GX-WT reactions. Each image is time-stamped in min:sec, and the sliding cargo MT plus ends are indicated by an asterisk. The angled sliding image series has two angled sliding events: one cargo MT slides only across the MT minus end of the template, and the other cargo MT initially slides across the lattice that transitions to across the template MT minus end. Scale bar, 10 μm. (C) Average MT sliding velocities of cargo MTs in anti-parallel or angled MT cross-links plotted as the mean ± SD and compared with a two-way ANOVA. Anti-parallel sliding velocities are the same as plotted in Figure 7C. Angled sliding events: GX-WT, n=60; GX-MBD1^mut^, n=30; GX-MBD2^mut^, n=40. N=4 independent experiments per condition. ns, not significant; *, p<0.05; **, p<0.01; ***, p<0.001; ****, p<0.0001.

**Video S1-3**. Aligned MT sliding of anti-parallel (left) and parallel (right) MT cross-links of GX-WT (S1), GX-MBD1^mut^ (S2), or GX-MBD2^mut^ (S3). Template MTs have magenta seeds and gray extensions, and cargo MTs have gray seeds and magenta extensions. Scale bar, 5 μm.

**Video S4**. Representative angled MT sliding events by GX-WT: angled (left) and aligning (right). Template MTs have magenta seeds and gray extensions, and cargo MTs have gray seeds and magenta extensions. Scale bar, 5 μm.

## Notes

### Competing Interest Statement

The authors have declared no competing interest.

### Summary of Updates

The new version includes additional analyses of the spindle assembly data (Figure 5) and the Microtubule Cross-linking experiments (Figures 6 and 7).

